# The neuroimmune CGRP-RAMP1 axis tunes cutaneous adaptive immunity to the microbiota

**DOI:** 10.1101/2023.12.26.573358

**Authors:** Warakorn Kulalert, Alexandria C. Wells, Verena M. Link, Ai Ing Lim, Nicolas Bouladoux, Motoyoshi Nagai, Oliver J. Harrison, Olena Kamenyeva, Juraj Kabat, Michel Enamorado, Isaac M. Chiu, Yasmine Belkaid

## Abstract

The somatosensory nervous system surveils external stimuli at barrier tissues, regulating innate immune cells under infection and inflammation. The roles of sensory neurons in controlling the adaptive immune system, and more specifically immunity to the microbiota, however, remain elusive. Here, we identified a novel mechanism for direct neuroimmune communication between commensal-specific T lymphocytes and somatosensory neurons mediated by the neuropeptide Calcitonin Gene-Related Peptide (CGRP) in the skin. Intravital imaging revealed that commensal-specific T cells are in close proximity to cutaneous nerve fibers *in vivo*. Correspondingly, we observed upregulation of the receptor for the neuropeptide CGRP, RAMP1, in CD8^+^ T lymphocytes induced by skin commensal colonization. Neuroimmune CGRP-RAMP1 signaling axis functions in commensal-specific T cells to constrain Type 17 responses and moderate the activation status of microbiota-reactive lymphocytes at homeostasis. As such, modulation of neuroimmune CGRP-RAMP1 signaling in commensal-specific T cells shapes the overall activation status of the skin epithelium, thereby impacting the outcome of responses to insults such as wounding. The ability of somatosensory neurons to control adaptive immunity to the microbiota via the CGRP-RAMP1 axis underscores the various layers of regulation and multisystem coordination required for optimal microbiota-reactive T cell functions under steady state and pathology.

**Significance statement:** Multisystem coordination at barrier surfaces is critical for optimal tissue functions and integrity, in response to microbial and environmental cues. In this study, we identified a novel neuroimmune crosstalk mechanism between the sensory nervous system and the adaptive immune response to the microbiota, mediated by the neuropeptide CGRP and its receptor RAMP1 on skin microbiota-induced T lymphocytes. The neuroimmune CGPR-RAMP1 axis constrains adaptive immunity to the microbiota and overall limits the activation status of the skin epithelium, impacting tissue responses to wounding. Our study opens the door to a new avenue to modulate adaptive immunity to the microbiota utilizing neuromodulators, allowing for a more integrative and tailored approach to harnessing microbiota-induced T cells to promote barrier tissue protection and repair.

## Introduction

Mammalian neural and immune cells are constantly exposed to external stimuli at barrier surfaces, and therefore have evolved to sense and respond to fluctuating environmental cues present in the tissue milieu. Correspondingly, to achieve such common sentinel goals, the nervous and immune systems communicate largely with shared signaling molecules such as cytokines, chemokines and neuropeptides, blurring the discrete functional partitioning between these two critical biological units (Talbot, Foster, and Woolf 2016; Chu, Artis, and Chiu 2020; Huh and Veiga-Fernandes 2020). Neuroimmune crosstalk, which can have multifaceted consequences on the barrier tissues, has thus emerged as a crucial research area that could result in a more holistic understanding of tissue physiology and pathology, as well as novel therapeutic avenues in the context of local infection and tissue inflammation (Ordovas-Montanes et al. 2015; Veiga-Fernandes and Mucida 2016; Klein Wolterink et al. 2022).

While neuroinflammation in neurodegeneration, in particular those involving the central nervous system and devastating neurological diseases in humans, have been well-characterized (Becher, Spath, and Goverman 2017; Kinney et al. 2018; Charabati et al. 2023), exhaustive examination of neuroimmune interaction in the peripheral tissues remains at its early stage. Of note, regulation of tissue immunity by the autonomic nervous system including sympathetic, parasympathetic or enteric neurons, in particular in the context of anti-inflammatory reflex, has been documented (Tracey 2002; Matheis et al. 2020). What remains to be further characterized are any direct immunological roles of somatosensory fibers, in particular the abundant nociceptive and/or peptidergic neuronal subsets (Le Pichon and Chesler 2014) that integrate external stimuli ranging from gentle touch to noxious pain at barrier sites. Our specific goal of this study is to examine how such nerve fibers, highly enriched in the skin, directly engage immune cells at homeostasis at this largest barrier site and organ.

Seminal studies have revealed that innate immune cells such as neutrophils and dendritic cells can be regulated by molecular mediators associated with the nervous system such as the neuropeptide CGRP in the context of bacterial and fungal infection (Chiu et al. 2013; Kashem et al. 2015; Pinho-Ribeiro et al. 2018; Lai et al. 2020) and skin inflammation (Riol-Blanco et al. 2014; Zhang et al. 2021), often involving pain sensation (Lagomarsino, Kostic, and Chiu 2021). Innate lymphoid cells, such as ILC2s in the lung and the intestine, have also been shown to respond to CGRP in the context of parasitic infection (Wallrapp et al. 2019; Xu et al. 2019; Nagashima et al. 2019). Substance P, another critical neuropeptide secreted by sensory fibers, can regulate immune cells involved in allergy such as mast cells (Serhan et al. 2019) and dendritic cells (Perner et al. 2020). Optogenetic activation of cutaneous nociceptors can elicit inflammation termed anticipatory Type 17 immune response in the skin (Cohen et al. 2019). What remains unclear is whether somatosensory neurons at barrier sites can directly control long-lasting and highly specific adaptive immune cells, particularly under homeostasis, in the absence of any infection or apparent tissue inflammation.

The skin is colonized by the microbiota, which in turn plays a fundamental role in educating and regulating host adaptive immune cells to maintain barrier tissue homeostasis (Honda and Littman 2016; Ansaldo, Farley, and Belkaid 2021). For instance, the human skin commensal *Staphylococcus epidermidis* robustly induces accumulation of T lymphocytes that harbor diverse functions beneficial to the mammalian host including tissue repair and antimicrobial defense under steady state, without any manifestations of inflammation or infection (Naik et al. 2015; Linehan et al. 2018; Harrison et al. 2019; Constantinides et al. 2019; Enamorado et al. 2023). Because these functionally unique and versatile commensal-specific T cells are situated in tissues that are highly innervated, it is crucial to understand the extent to which microbiota-reactive T cells interact with and are controlled by somatosensory nerves in the barrier tissues. In this study, we identified a novel mechanism by which sensory neurons directly regulate commensal-reactive T cells via the neuropeptide CGRP and its cognate receptor RAMP1 on T cells in the skin. The neuroimmune CGRP-RAMP1 axis functionally tunes commensal-specific T cells to constrain Type 17 responses and control T cell activation status under steady state, with functional consequences upon overall tissue activation and responses to insults, demonstrating the direct regulation of homeostatic adaptive immunity by sensory neurons, shaping barrier tissue physiology in the milieu enriched with a diverse array of microbes and sensory modalities.

## Results

### Commensal-specific T cells are in close proximity to sensory nerve fibers in the skin at homeostasis

To assess cutaneous neuroimmune crosstalk, we first examined if there was a physical association between commensal-reactive T lymphocytes and somatosensory fibers innervating the skin. To this end, the skin of specific-pathogen free mice was colonized topically with a new commensal microbe, *Staphylococcus epidermidis*, inducing an accumulation of commensal-reactive CD8^+^ T cells without any manifestations of inflammation (Naik et al. 2012; Naik et al. 2015). After topical association with the commensal, we employed confocal microscopy to visualize microbiota-reactive, polyclonal CD8^+^ T cells that accumulated in the mouse skin, and observed that these T cells were in close proximity to sensory nerve peripherals, with some in direct physical contact with the sensory fibers (**Figure 1A**), suggesting a potential functional neuroimmune interaction at this barrier tissue under steady state.

**Figure 1.**
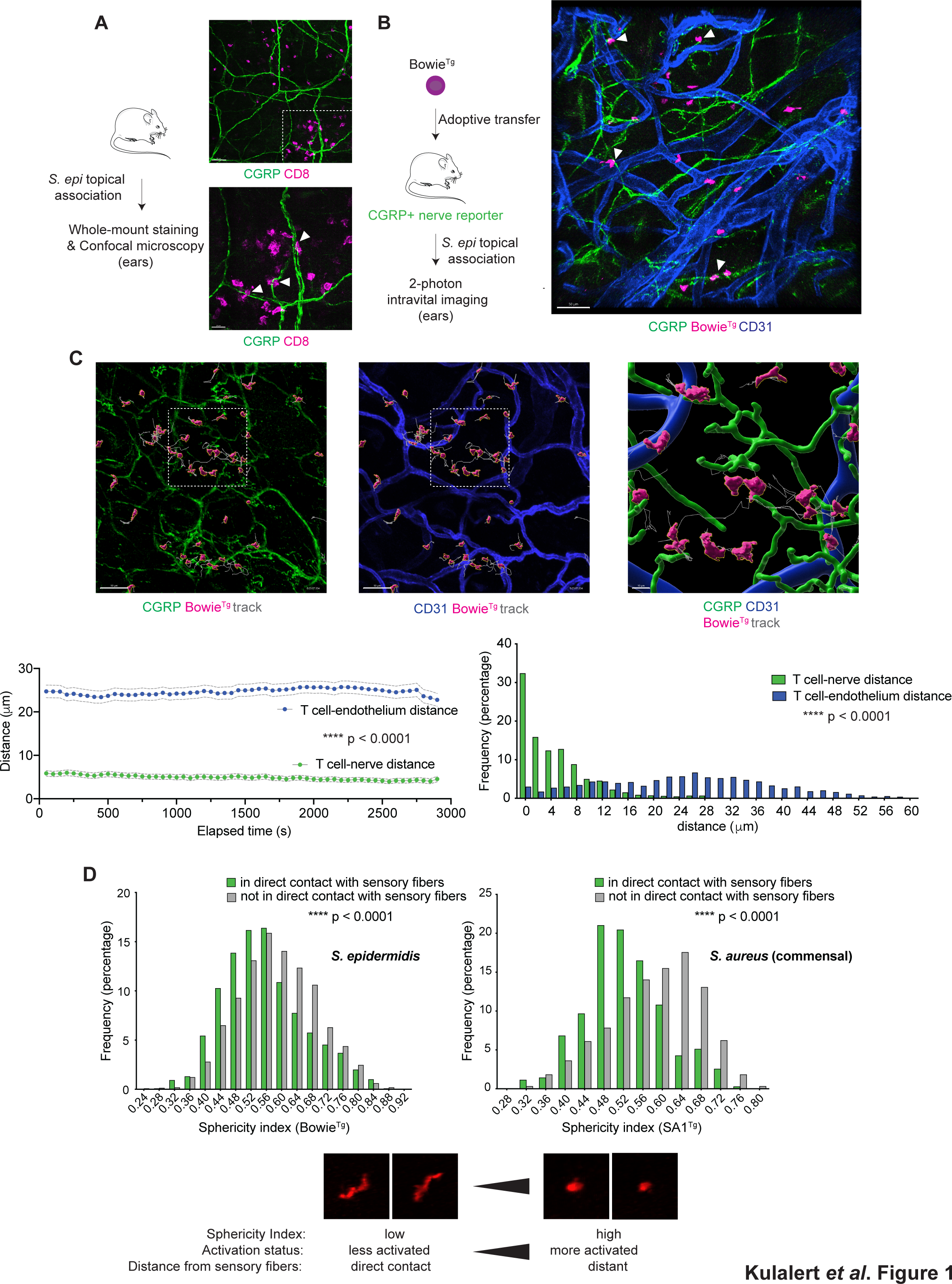
Commensal-specific T lymphocytes are in close proximity to sensory nerve fibers in the skin *in vivo* **(A)** Specific pathogen-free (SPF) C57BL/6 mice were topically associated with *Staphylococcus epidermidis* isolate LM087 in Tryptic Soy Broth (TSB) every other day for four times. Fourteen days after the initial association, mouse ears were fixed and stained for CGRP+ sensory neurons (anti-CGRP isoform α; green) and CD8^+^ T cells (anti-CD8α; magenta). Unassociated ears do not have CD8^+^ T cell accumulation, and therefore CD8^+^ T cells accumulating in the skin shown were induced by new commensal colonization. The bottom panel depicts magnification of a region demarcated by white dash lines in the top panel. White scale bars represent 50μm (bottom left corner of top panel) and 20μm (bottom left corner of bottom panel). Arrowheads indicate direct contact between T cells and nerve fibers. **(B)** Adoptive transfer of OFP^+^ *S. epidermidis*-specific transgenic T cells (Bowie^Tg^; magenta) was performed on albino SPF C57BL/6 hosts harboring the *Calca-EGFP* reporter (marking CGRP^+^ sensory fibers; green) prior to intravital imaging. Anti-CD31 (marking the vasculature; blue) injection was performed prior to imaging. A white scale bar (bottom left) represents 50μm. Arrowheads denote direct contact between commensal- specific CD8^+^ T cells and CGRP^+^ sensory fibers. **(C)** Each image panel represents the same visual field, with the same experimental approach and color schematic described in (**B**). Note the gray track representing dynamic *in vivo* T cell motility/displacement over the recording period, shown in **Supplemental Video 1**. The 3D cartoon rendering in the rightmost panel represents magnification of the region demarcated by white dashed lines in the leftmost and middle panels. Graphs represent quantitation of shortest distances between the T cells and the indicated neural or endothelial structures. Each data point in the left graph represents an average distance between T cells and respective structures at each elapsed time point, with standard deviation shown in dashed lines. The right graph is a histogram, illustrating the distribution of distances between T cells and respective structures over the recording period. **(D)** Each graph represents distribution of cellular sphericity indexes of commensal-specific T cells (Bowie^Tg^ for *S. epidermidis* and SA1^Tg^ for *S. aureus*) over the recording period. Bars in the indicated colors represent populations of the T cells that are in direct contact or not in direct contact with sensory fibers. The bottom schematic shows representative images of the cells with the indicated range of sphericity values and summarizes the relationship between distances from nerve fibers and sphericity and potential activation status. All data shown in (**A**)-(**D**) represent at least three independent experiments. ****p < 0.0001 as calculated with Student’s t test. See also **Figure S1**.

To further study the dynamic interaction between commensal-specific T cells and sensory nerve fibers *in vivo*, we utilized intravital 2-photon microscopy. Mice carrying a reporter for sensory fibers (*Calca-EGFP*, McCoy et al., 2012) were adoptively transferred with TCR transgenic *S. epidermidis*-specific CD8^+^ T cells (Bowie^Tg^) and topically associated with the commensal. Following topical association, Bowie^Tg^ T cells were in close proximity to sensory nerves, with an average distance of 4.9 ± 5.6 μm over a defined intravital recording period *in vivo*. Notably, a substantial portion (over 32%) of the Bowie^Tg^ cells were in direct contact (0 μm) with the neural fibers (**Figures 1B** and **1C**; **Supplemental Video 1**). In contrast, the Bowie^Tg^ T cells were comparatively distant to the vasculature over the same intravital recording period, with an average distance 24.6 ± 13.2 μm (**Figure 1C**). Overall, the Bowie^Tg^ T cells were evenly distributed spatially in relation to the endothelia over the *in vivo* recording period (low skewness of 0.1, with median distance of 25 μm), in contrast with the T cell-nerve distance distribution that was skewed towards direct contact (high skewness of 1.7, with median distance of 3.25 μm) over the recording period (**Figure 1C**). Of note, we previously reported a significant physical interaction between commensal-specific T cells and somatosensory nerve peripherals in CD4^+^ T cells induced by another skin commensal bacterium, *Staphylococcus aureus* (Enamorado et al. 2023), suggesting that neuroimmune interaction may be an evolutionarily conserved feature for conventional αβ T cells induced by a broad array of commensal microbes at the barrier site under steady state.

T cell shape and motility have been associated with antigen-mediated signal strength (Moreau et al. 2015) and activation status (Negulescu et al. 1996). We thus asked whether physical proximity to sensory nerve fibers could impact commensal-specific T cell behavior *in vivo*. Following *S. epidermidis* association, T cells that were in direct contact with the sensory fibers (distance = 0 μm) exhibited lower sphericity in morphology than those situated further from the nerves (distances > 0 μm) over the intravital recording period (**Figure 1D**, left panel). Our observations point to the potential ability of sensory fibers to influence T cell functions, as T cells that are distant from the sensory fibers tend to manifest rounding, associated with heightened activation status (Negulescu et al. 1996). This correlation between physical proximity and cellular shape *in vivo*, while prominent in the context of the nerve-T cell interface, was not evident in the context of the vasculature-T cell interface (**Figure S1**). Furthermore, this unique *in vivo* relationship between T cell-nerve interaction and cellular shape was also observed in commensal-specific CD4^+^ T cells upon topical colonization by another commensal *Staphylococcus* species, *S. aureus* (**Figure 1D**, right panel) under steady state, indicating that such manifestation of neuroimmune coordination may be an evolutionarily conserved feature in the context of homeostatic immune response to a broad range of commensals, and may confer functional consequences in the tissue under homeostasis.

Taken together, these observations of intimate physical neuroimmune interaction motivated us to further address whether the sensory nervous system could directly regulate commensal-induced lymphocytes.

### Skin commensal-induced T cells upregulate RAMP1, the receptor for neuropeptide CGRP

Our observations that commensal-specific T cells were closely associated with sensory nerve peripherals raised mechanistic questions underlying the neuroimmune crosstalk in the skin. We thus examined whether commensal-specific T cells expressed any molecular machinery that equipped them with the potential ability to respond to neuronally derived signals such as neuropeptides. Unbiased transcriptomic analysis revealed that a *S. epidermidis*-induced population of CD8^+^ T cells (Tc17: CCR6^+^ CD8^+^) exhibited a significant transcriptional upregulation of the *Ramp1* gene encoding the receptor for neuropeptide CGRP (Figure 2A, RNA-seq dataset from our laboratory, Linehan et al., 2018). In contrast, naïve and antigen- experienced (T_EM_) CD8^+^ T cells isolated from the skin-draining lymph nodes of commensal- colonized animals did not exhibit any significant induction of *Ramp1* (**Figure 2A**). Of note, in our examination of over 100 neuronal genes expressed in diverse skin CD8^+^ T cell populations, we observed upregulation of a number of neuropeptide, neurotrophic factor or neuroendocrine signaling-related genes (*Ramp1, Ramp3, Ntrk3, Calca, Gch1, Nr3c1* and *Bex3*) in commensal-induced Tc17 cells, compared to CD8^+^ T cells induced under infection. These data suggest a wide range of signaling molecules that could be associated with neuronal control of adaptive immunity to commensals, with *Ramp1* being the most induced neuroimmune mediator gene candidate (**Figure S2A**). Upregulation of *Ramp1* was unique to commensal-induced Tc17 cells, as *Ramp1* expression was modest or not detected in pathogenic CD8^+^ T cells accumulating in the skin after *S. epidermidis* intradermal infection, HSV infection or *Leishmania* infection (**Figures 2B** and **S2B**). Expression of another CGRP co-receptor encoding gene *Calcrl* remained low and undistinguishable across the skin T cell populations under diverse conditions (**Figure S2C**), suggesting that dynamic expression of *Ramp1* likely plays a dominant role in CGRP signaling in T cells, at least in the host-microbiota dialog in the skin compartment.

**Figure 2.**
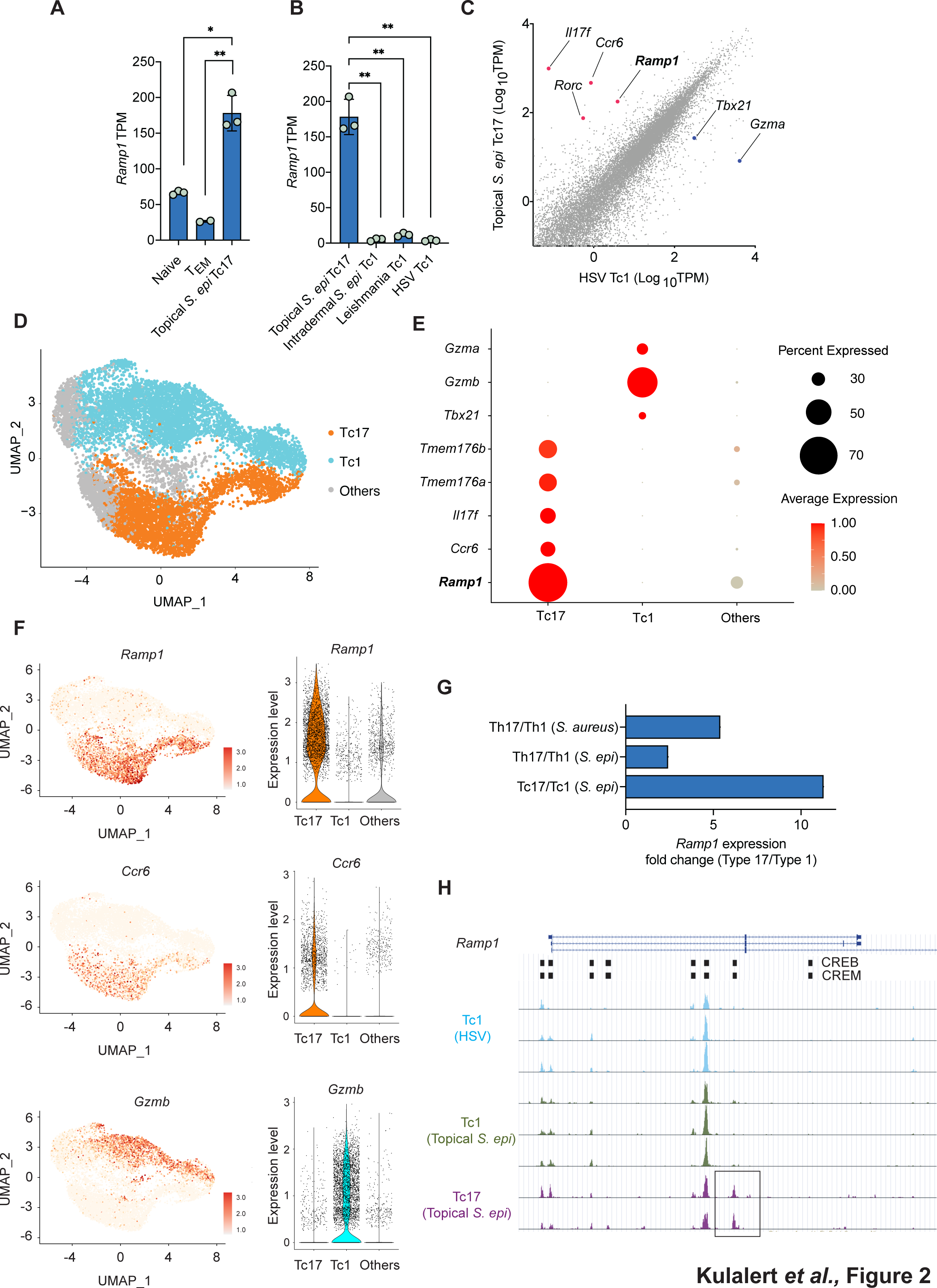
Skin microbiota-reactive T cells express the receptor for the neuropeptide CGRP at homeostasis *in vivo* **(A)** Comparison of *Ramp1* expression levels among the indicated T cell populations. “Naïve” refers to naïve T cells isolated from the skin-draining lymph nodes of *S. epidermidis*-associated animals. “T_EM_” refers to effector memory T cells isolated from the skin-draining lymph nodes of *S. epidermidis*-associated animals. Tc17 refers to CCR6^+^ CD8^+^ T cells. **(B)** Comparison of *Ramp1* expression levels among the indicated T cell populations from the skin. **(C)** Scatter plot showing differentially expressed genes, comparing topical *S. epidermis*- induced Tc17 cells vs HSV-induced Tc1 (CCR6^-^ CD8^+^) cells from the skin. Red and blue large dots highlight differentially expressed genes of interest. **(D)** Single-cell RNA-seq data of sorted skin CD8^+^ T cells from animals topically associated with *S. epidermidis*. Clusters were assigned based on expression levels of canonical Tc1 and Tc17 markers, including those shown in (**E**). **(E)** Dot plot for scRNA-seq data, showing the frequency of each population of skin CD8^+^ T cells expressing the indicated Tc1 or Tc17 markers and *Ramp1*, as well as the expression level for each transcript. Dots are colored by average expression and scaled by number of cells expressing the indicated transcript within each cluster (frequency). **(F)** Feature plots and violin plots for scRNA-seq data, showing the distribution and the frequency of each population of skin CD8^+^ T cells expressing key transcripts indicated. **(G)** Comparison of expression of *Ramp1* in microbiota-induced Type 17 T cells vs Type 1 T cells elicited by the indicated commensal colonization from RNA-seq datasets generated in our laboratory. **(H)** UCSC Genome Browser image of open chromatin regions of the *Ramp1* locus in the indicated CD8^+^ T cell populations from the ATAC-seq dataset. Refer to detailed RNA-seq and ATAC-seq data sources and experimental settings in **Materials and Methods**. Bar graphs show mean ± standard deviation. P-values were calculated with Student’s t test. See also **Figure S2**.

*Ramp1* was significantly upregulated in commensal-induced Tc17 cells in comparison to another commensal-induced population, Tc1 cells (CCR6^-^ CD8^+^) (**Figure S2D**). Furthermore, the magnitude of *Ramp1* upregulation in commensal-induced Tc17 vs *Ramp1^low^* Tc1 cells was substantial, comparable to the induction of key Type 17 response genes such as *Ccr6, Il17f* and *Rorc* in all Tc17 vs Tc1 comparisons performed under different conditions (topical *S. epi* vs HSV in **Figure 2C**, topical vs intradermal *S. epi* in **Figure S2B** and Tc17 vs Tc1 in **Figure S2D**). Single-cell transcriptomic analysis of CD8^+^ T cells isolated from the skin following *S. epidermidis* topical association (**Figure 2D**, scRNA-seq dataset from our laboratory, Harrison et al. 2019) further corroborated that *Ramp1* upregulation was enriched in the Tc17 population (defined as the cluster with induction of Type 17 response genes including *Ccr6, Il17a, Il17f, Tmem176a, Tmem176b*), in contrast with the Tc1 population (defined as the cluster with induction of Type 1 response genes including *Tbx21, Gzma, Gzmb*) (**Figures 2E** and **2F**).

Additionally, we observed upregulation of *Ramp1* in *S. epidermidis*-induced Type 17 CD4^+^ T cells (Th17), which also accumulate in the skin after *S. epidermidis* topical colonization (Naik et al. 2015), compared to their *Ramp1^low^*Th1 counterparts (**Figure S2E**). The magnitude of *Ramp1* upregulation in Th17 was comparable to key Type 17 response genes such as *Ccr6, Il17f* and *Rorc* (**Figure S2E**). Furthermore, elevated expression of *Ramp1* was observed in Th17 cells accumulating in the skin after topical association with another skin commensal, *S. aureus* (Enamorado et al. 2023). Of note, mucosal-associated invariant T cells, which are imprinted by the microbiota (Constantinides et al. 2019), also exhibited upregulation of *Ramp1* in their CCR6^+^/Type 17 subsets in various barrier compartments (**Figure S2F**, scRNA-seq dataset from our laboratory, Constantinides et al. 2019). Taken together, these observations illustrate that upregulation of the receptor for the neuropeptide CGRP, RAMP1, is a conserved attribute of diverse populations of Type 17 classical (**Figure 2G**) and non-classical (**Figure S2F**) T cells and may contribute to neuroimmune crosstalk in response to a broad array of commensal microbes at barrier surfaces under homeostasis.

To further understand the mechanisms underlying upregulation of the neuroimmune CGRP- RAMP1 axis in commensal-induced T cells under steady state, we analyzed ATAC-seq data, previously generated in our laboratory (Harrison et al. 2019), performed in Tc1, Tc17, Th1 and Th17 cells accumulating in the skin following *S. epidermidis* topical association. Notably, we identified regions of accessible chromatin corresponding to regulatory elements of *Ramp1* that are unique to commensal-induced Tc17 and Th17 populations, compared to their Tc1 and Th1 counterparts (boxed regions in **Figures 2H** and **S2G**), consistent with transcriptional upregulation of *Ramp1*. Notably, these open chromatin regions of *Ramp1* enriched in Tc17 and Th17 cells contained cAMP-responsive elements such as CREB and CREM motifs (**Figure 2H**), pointing to potential engagement of Type 17 cell-specific *Ramp1* enhancers with cAMP and its transducers known to be downstream of CGRP signaling (Harzenetter et al. 2007; Russell et al. 2014). These observations may mechanistically account for transcriptional upregulation of *Ramp1* in commensal-induced skin Tc17 and Th17 populations in at homeostasis, pointing to a role for CGRP signaling in reinforcing positive feedback on its own receptor expression via cAMP-dependent transcription factors that can bind to Type 17 cell-specific open chromatin regions of *Ramp1*.

Collectively, our observations unveiled a robust induction of the CGRP receptor RAMP1 specifically in Type 17 commensal-induced T cells, potentially endowing the T cells with an ability to communicate directly with the sensory nervous system in the skin under steady state.

### Neuroimmune CGRP-RAMP1 signaling constrains Type 17 responses in commensal-induced T cells

To examine the extent to which CGRP-RAMP1 signaling in commensal-reactive T cells impacts lymphocyte functions, we genetically ablated *Ramp1* expression specifically in the T cells, utilizing a conditional knockout model, breeding *Ramp1 fl/fl* mice with *Lck Cre* mice. Because CGRP-RAMP1 signaling functions in many compartments in the skin including sensory nerves, the endothelium and innate immune cells (Russell et al. 2014), a T cell-specific RAMP1 depletion approach is essential to unambiguously exclude any indirect or cell-nonautonomous impacts of CGRP on commensal-induced T cells at the barrier site.

To characterize the mechanisms underlying neuroimmune CGRP-RAMP1 signaling in commensal-induced T cells under steady state in an unbiased manner, we performed single-cell RNA-sequencing on CD8^+^ T cells isolated from the skin of mice lacking *Ramp1* expression specifically in the T cell compartment (*Lck Cre+ Ramp1 fl/*fl) vs control animals (*Lck Cre- Ramp1 fl/fl*), following *S. epidermidis* topical colonization (**Figure 3A**). Consistent with the RNA-seq data described in **Figure 2**, we observed upregulated *Ramp1* expression exclusively in the Tc17 populations, but not in the Tc1 or other populations (**Figures 3B, 3C** and **3D**). Importantly, the *Ramp1* expression level in these Tc17 cells was significantly attenuated in the *Lck Cre+ Ramp1 fl/fl* genetic background compared to control (**Figures 3C** and **3D**), confirming our robust T cell-specific *Ramp1* ablation approach. Of note, the number and the frequency of cells in each of the dominant Tc17 clusters (clusters 0 and 10) in the T cell-specific *Ramp1*- deficient genetic background were increased, compared to control (**Figures 3E** and **S3A**), indicating that RAMP1 may act to constrain accumulation and/or activation of commensal-induced Tc17 cells in the skin at homeostasis.

**Figure 3.**
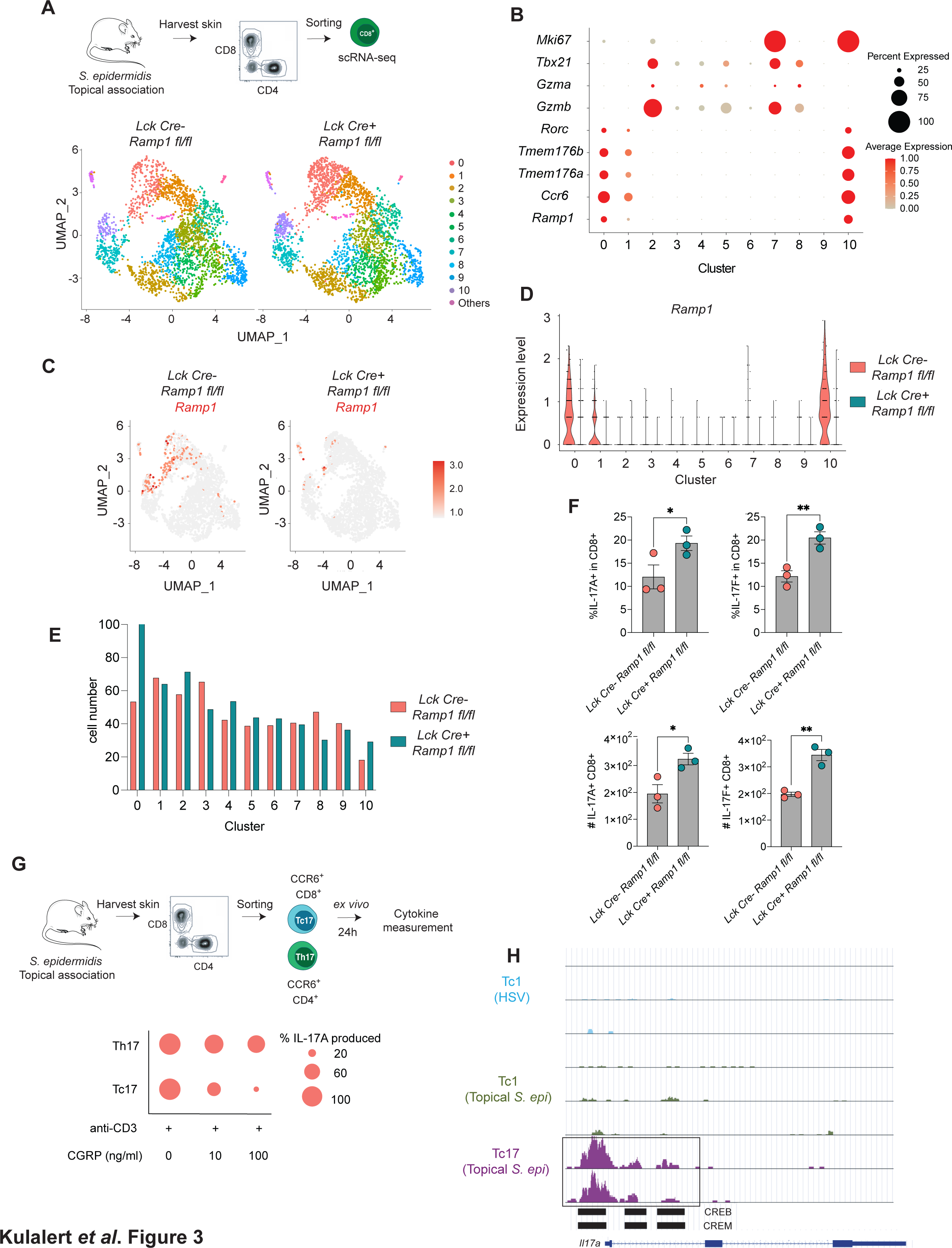
The neuroimmune CGRP-RAMP1 axis constrains Type 17 responses in commensal-induced T cells *in vivo* **(A)** A schematic for the scRNA-seq experimental workflow (top). UMAP plots show all sorted CD8^+^ T cells from *Ramp1*-deficient and control mice (bottom). Assigned Seurat clusters are numbered and color-coded. **(B)** Dot plot showing the frequency of each population of skin CD8^+^ T cells expressing the indicated Tc1 or Tc17 markers and *Ramp1*, as well as the expression level for each transcript, from both *Ramp1*-deficient and control mice combined. Dots are colored by average expression and scaled by number of cells expressing the indicated marker within each cluster (frequency). **(C)** UMAP plots showing *Ramp1* expression levels in all cells for each indicated genotype. **(D)** Violin plot showing expression of *Ramp1* for each Seurat cluster for each indicated genotype. **(E)** Bar graph representing the cell number from each genotype (*Ramp1*-deficient vs control genetic background) for each assigned Seurat cluster. **(F)** Bar graphs illustrating flow cytometry data from each indicated genotype. Data represent two independent experiments. Bar graphs show mean ± standard error of the mean. P- values were calculated with Student’s t test. **(G)** Dot plot (bottom) showing the extent to which IL-17A was produced by Tc17 and Th17 cells isolated and treated *ex vivo* based on the schematic shown (top). IL-17A production levels are shown in percentages, relative to control (with anti-CD3e, no CGRP), as reflected by the scaling of each dot. Data represent two independent experiments. **(H)** UCSC Genome Browser image of open chromatin regions of the *Il17a* locus in the indicated CD8^+^ T cell populations in the wild-type genetic background from the ATAC- seq data described in Figure 2. See also **Figure S3**.

To determine whether the CGRP-RAMP1 axis modulates Type 17 responses in commensal-induced T cells, we examined the skin immune cells of animals lacking *Ramp1* in the T cell compartment vs control animals by flow cytometry following *S. epidermidis* topical association. We found that RAMP1 deficiency in T cells significantly enhanced both the frequency and the cell number of the skin commensal-induced CD8^+^ T cells that produced IL-17A and IL-17F, compared to control, under steady state in the skin (**Figure 3F**). Consistent with RAMP1 functioning to restrain Type 17 responses in commensal-induced T cells, we observed an overall upregulation of key Type 17 mRNA transcripts in the T cell-specific *Ramp1*-deficient CD8^+^ T cells, compared to wild-type control under steady state (**Figure S3B**). To determine whether the neuropeptide CGRP can regulate Type 17 cytokine production by T cells, we then utilized an *ex vivo* culture approach, assessing IL-17A production in sorted and cultured commensal-induced Tc17 and Th17 cells in the presence of different concentrations of CGRP with TCR stimulation via administration of anti-CD3e (**Figure 3G**). Commensal-induced Tc17 and Th17 cells isolated from the skin exhibited diminished IL-17A production in response to CGRP in a dose-dependent manner *ex vivo* (**Figure 3G**). Taken together, these observations revealed a novel neuroimmune role of CGRP, demonstrating that T cell-intrinsic CGRP-RAMP1 signaling functions at the barrier site to constrain Type 17 responses in microbiota-induced T lymphocytes at homeostasis.

To gain further insights into how CGRP-RAMP1 signaling controls Type 17 T cell responses to the microbiota under steady state, we utilized our previously generated ATAC-seq (Harrison et al. 2019). With this approach, we identified regions of open chromatin unique to Tc17 and Th17 cells in Type 17 response genes (boxed regions in **Figures 3H, S3C** and **S3D**), in the wild-type genetic background. Importantly, we discovered cAMP-responsive elements such as CREB and CREM motifs (boxed regions in **Figures 3H, S3C** and **S3D**) in the Tc17/Th17-specific open chromatin regions of Type 17 response genes. Our motif discovery suggests potential transcriptional regulation of Type 17 gene expression by cAMP and its transducers, downstream of neuroimmune CGRP-RAMP1 signaling, to constrain Type 17 responses in commensal-induced T cells in the skin at homeostasis.

### Neuroimmune CGRP-RAMP1 signaling tunes T cell activation status and shapes barrier tissue physiology

The impact of CGRP-RAMP1 signaling on Type 17 responses in T cells suggested that T cell behavior and activation status may also be subject to neuroimmune control in the skin. To address this point, we generated transgenic commensal-specific T lymphocytes (Bowie^Tg^) that can be distinguished visually between wild-type (OFP^+^) and *Ramp1*-deficient (GFP^+^) upon adoptive transfer to a common host and tissue environment (**Figure 4A**). Following *S. epidermidis* colonization, *Ramp1*-deficient Bowie^Tg^ T cells in the skin exhibited higher sphericity in their morphology over the intravital recording period compared to wild-type Bowie^Tg^ T cells (**Figure 4A**), indicating that the T cells without CGRP-RAMP1 signaling may have heightened activation status, as high sphericity/rounding is associated with elevated calcium levels in T cells (Negulescu et al. 1996). Indeed, the enhanced sphericity in *Ramp1*-deficient Bowie^Tg^ T cells was reminiscent of rounding observed in the population of T cells that were distant from sensory fibers, as described in **Figure 1**. Of note, *Ramp1*-deficient Bowie^Tg^ T cells overall exhibited a similar morphology pattern to the population of wild-type Bowie^Tg^ T cells that were distant from sensory nerve fibers (higher sphericity *in vivo*, **Figure 1D**), pointing to a convergent state of heightened T cell activation/rounding that occurs when neuroimmune communication is disrupted or attenuated by either increased physical distances (**Figure 1D**) or genetic ablation of *Ramp1* (**Figure 4A**). Furthermore, the relationship between T cell morphology and physical association with sensory fibers *in vivo* was dominantly RAMP1-dependent (**Figure S4A**). Taken together, our intravital neuroimmune observations propose that the neuroimmune CGRP-RAMP1 axis modulates cellular morphology and behavior, potentially reflecting commensal-specific T cell activation status in the skin.

**Figure 4.**
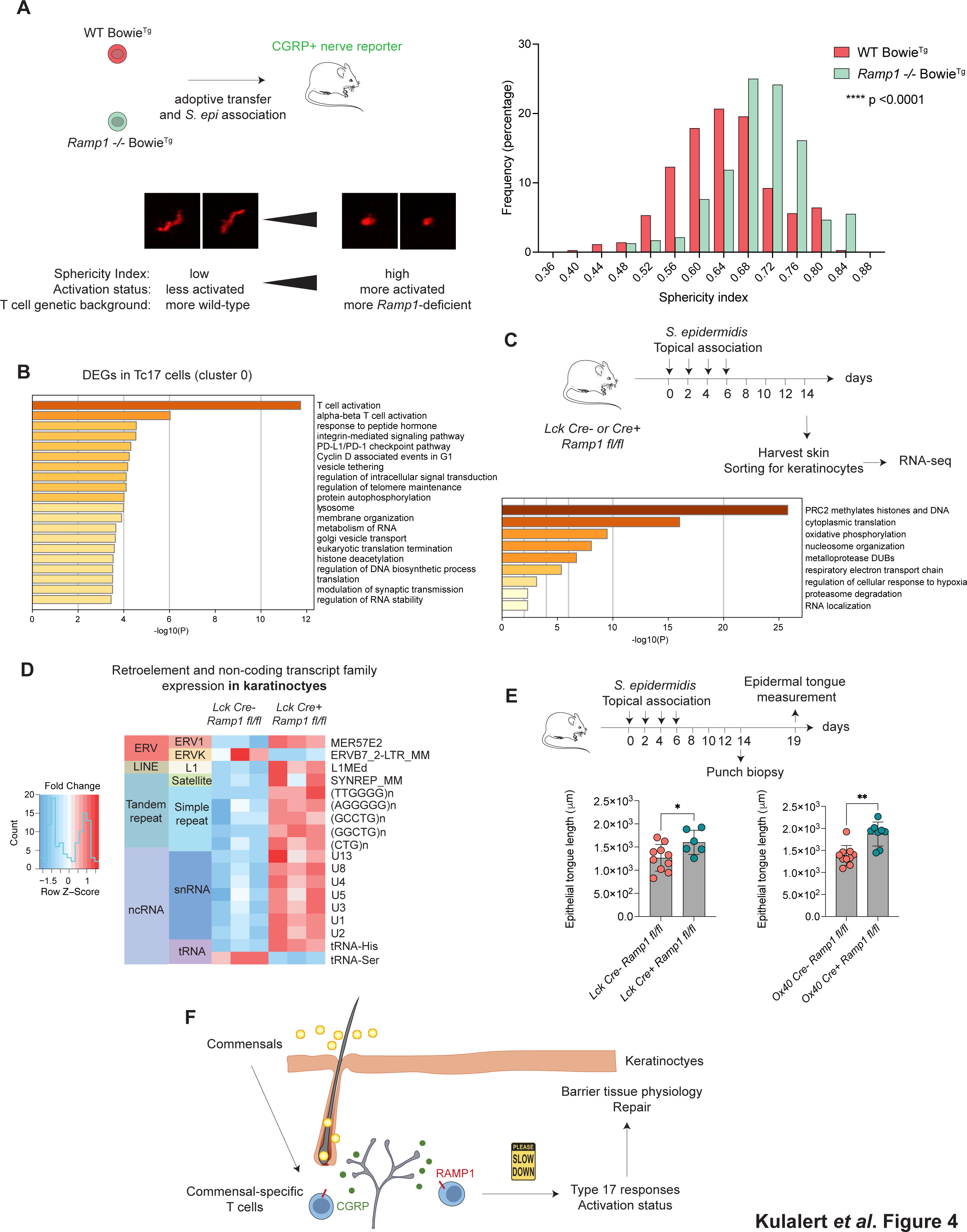
Neuroimmune CGRP-RAMP1-mediated regulation of microbiota-reactive T cells shapes barrier tissue physiology **(A)** The top schematic depicts the experimental setup where commensal-specific (Bowie^Tg^) T cells from *Ramp1*-deficient and control genetic backgrounds, expressing distinct fluorophores, were adoptively transferred to common hosts harboring the *Calca-EGFP* reporter for sensory fiber visualization, followed by *S. epidermidis* topical association. The graph represents distribution of cellular sphericity values over the recording period for each indicated genotype of the commensal-specific T cells. P-values were calculated with Student’s t test. The bottom schematic shows representative images of the cells with the indicated range of sphericity values and summarizes the relationship between cell shape/potential activation status and *Ramp1* genetic background. Data represent two independent experiments. **(B)** GO enrichment analysis of differentially expressed genes in the dominant, non- proliferating Tc17 cluster (Cluster 0 from the scRNA-seq dataset described in Figure 3), comparing commensal-induced T cells from *Ramp1*-deficient vs control genetic backgrounds. **(C)** GO enrichment analysis of differentially expressed genes, comparing keratinocytes from the skin milieu where commensal-induced T cells were *Ramp1*-deficient vs control. **(D)** Analysis of families of differentially expressed retroelement and non-coding transcript loci, comparing keratinocytes from the skin milieu where commensal-induced T cells were *Ramp1*-deficient vs control. **(E)** Quantification of the epidermal tongue length five days after wounding in male animals from the indicated genotypes, following *S. epidermidis* topical association (full body). Bar graphs show mean ± standard deviation. P-values were calculated with Student’s t test. Data represent three independent experiments. **(F)** A schematic summarizing the neuroimmune CGRP-RAMP1 axis functioning to constrain commensal-induced T cells and regulate barrier tissue physiology. See also **Figures S4** and **S5**.

To assess if T cell activation status was altered in the absence of CGRP-RAMP1 signaling, we compared transcriptomic profiles between wild-type and *Ramp1*-deficient commensal-induced T cells in our scRNA-seq dataset described in **Figure 3**. Because *Ramp1* is expressed and acts mainly in Tc17 cells (**Figures 2** and **3**), we focused on transcriptomic differences (*Lck Cre-* vs *Lck Cre+*) in two dominant Tc17 populations—clusters 0 and 10, all with high baseline *Ramp1* expression in the *Cre-* control background vs efficient *Ramp1* ablation in the *Cre+* background. Of note, the most striking transcriptomic distinction between wild-type and *Ramp1*- deficient commensal-induced Tc17 cells was observed in the most abundant Tc17 cluster (cluster 0), where the differentially expressed genes exhibited highly significant enrichment in factors controlling T cell activation (**Figure 4B**). Notably, in this dominant Tc17 cluster, we observed upregulation of genes encoding T cell activation markers TIM-3, PD-1 and CD44 in *Ramp1*- deficient T cells (**Figure S5A**), consistent with an overall increase in activation in Tc17 cells, as reflected in heightened Type 17 responses (**Figure 3**) and altered cellular morphology (**Figure 4A**). Another Tc17 cluster (cluster 10; *Mki67^high^*) also harbored *Ramp1*-dependent differentially expressed genes, enriched in biological processes involving cell division, indicating that the CGRP-RAMP1 axis may also control proliferation in this actively dividing subset of Tc17 cells (**Figures S5B** and **S5C**). Collectively, these observations point to a novel role of neuroimmune CGRP-RAMP1 signaling in regulating T cell activation status, specifically in commensal-induced Tc17 cells at the barrier site at homeostasis.

To assess the functional significance of neuroimmune-mediated constraint of microbiota-reactive T cells in the skin, we asked if heightened activity of *Ramp1*-deficient T cells may impact the broader activation status and physiology of the skin.

To this end, we focused on keratinocytes, which play a major role in maintaining cutaneous tissue integrity in the host-microbiota dialog (Naik et al. 2015). Specifically, we compared the transcriptional profiles of keratinocytes in the tissue environment where bystander commensal-reactive T cells were either *Ramp1*-deficient (*Lck Cre+ Ramp1 fl/fl*) or wild-type control (*Lck Cre- Ramp1 fl/fl*). Of note, our transcriptomic analysis showed that keratinocytes did not express *Ramp1*, and therefore any alterations in the keratinocyte transcriptomes would be attributed to cell-nonautonomous sources such as differential *Ramp1* expression levels in the T cell compartment. We observed that keratinocytes in the milieu with *Ramp1*-deficient commensal-induced T cells exhibited upregulation of genes associated with ectopic tissue activation, such as those involved in histone modification, nucleosome organization, electron transport chain and protein translation (**Figure 4C**), corresponding to the heightened activation status of bystander *Ramp1*-deficient T cells under steady state. Furthermore, we also observed upregulation of defined retroelements and non-coding transcripts in keratinocytes from animals with *Ramp1*- deficient commensal-induced T cells (**Figures 4D** and **S5D**), which have been associated with increased immune response or inflammation in defined settings (Lima-Junior et al. 2021).

To further assess whether the control of Type 17 responses and keratinocyte activation status by neuroimmune CGRP-RAMP1 signaling can confer functional consequences on the barrier tissue *in vivo*, we assessed epithelial wound repair, which we had previously shown to be linked to commensal-induced T cells and Type 17 responses in the skin (Linehan et al. 2018; Konieczny et al. 2022). We observed that the lack of neuroimmune CGRP-RAMP1 signaling axis in commensal-induced T cells resulted in enhanced wound healing upon injury (**Figure 4E**, also confirmed with an independent activated T cell-specific *Ox40 Cre* system), in accordance with the heightened Type 17 responses in *Ramp1*-deficient T cells. The impact of the CGRP-RAMP1 axis on tissue repair demonstrates that alterations in neuroimmune signaling in commensal-induced T cells enhance Type 17 responses and the overall activation status of keratinocytes, thereby shaping the outcome of biological processes governing tissue health and integrity.

Taken together, our observations illustrate that neuroimmune CGRP-RAMP1 signaling in microbiota-reactive T lymphocytes constrains Type 17 response and controls T cell activation, shaping the tissue milieu and profoundly impacting the activation status of the epithelial compartment (**Figure 4F**), which can have vast implications on tissue physiology particularly under additional insults or pathology.

## Discussion

In this study, we have established and delineated the physical and functional interface between the sensory nervous system and adaptive immunity to the microbiota at the largest barrier tissue at homeostasis. First, we demonstrated close physical proximity between commensal-specific T lymphocytes and cutaneous nerve peripherals. We then identified a key neuropeptide and its receptor (CGRP-RAMP1) as a novel signaling mechanism mediating the direct communication between local neural fibers and microbiota-reactive T cells in the skin. We then proceeded to elucidate the mechanisms by which the neuroimmune CGRP-RAMP1 signaling axis modulated Type 17 responses and shaped the transcriptome and activation status of commensal-induced Type 17 T cells under steady state, contributing to remodeling of the barrier site milieu. Collectively, these observations unveil a novel role of CGRP as a physiological neuroimmune signaling molecule that can impact adaptive immunity and subsequently the skin tissue environment and barrier integrity.

Previous studies have established neuroimmune communication in the context of innate immunity under infection or tissue inflammation in diverse barrier tissues in mammals (Chu, Artis, and Chiu 2020), as well as in invertebrate model organisms of host-microbe interactions (Makhijani et al. 2017; Kim and Flavell 2020). Our study has further identified a novel neuroimmune regulation mechanism in which adaptive lymphocytes can directly respond to neuronal signals under steady state, in the absence of pathogens or pro-inflammatory cues. Specifically, under homeostasis, commensal-specific T cells, which are highly plastic and can promote antimicrobial defense and repair, are capable of directly communicating with sensory fibers via CGRP. The ability of these lymphocytes to respond to constantly fluctuating sensory signals relayed to them by sensory neurons via CGRP is another testament to multisystem coordination and adaptability that the host has evolved to maintain barrier site homeostasis, particularly in the context of a complex host-microbiota dialog. Indeed, commensal-reactive T cells expressed additional neuropeptide, neurotrophin or neuroendocrine signaling-related genes in addition to those involved in the CGRP-RAMP1 signaling axis, indicating that there are a variety of neuroimmune signal transduction pathways beyond CGRP-RAMP1 to tune adaptive immunity to the microbiota. Of note, catecholamines and adrenergic signaling have been shown to regulate CD8^+^ T cell effector functions in the context of viral infection and anti-tumor therapy (Globig et al. 2023). Furthermore, we found that chemokine receptor genes *Ccr4* and *Tnfsf11* were induced in commensal-induced Tc17 cells in the skin, compared to pathogenic Tc1 cells (data not shown). Notably, sensory neurons express chemokines (Chiu et al. 2014), and therefore corresponding receptor expression in T cells represents yet another neuroimmune communication mechanism in parallel to the CGRP-RAMP1 axis. Collectively, these diverse neuroimmune molecules expressed at the T cell-nerve interface highlight a dynamic and multipronged dialog between the immune system and the sensory nervous system. Of note, sensory neurons have been shown to exert multimodal control on another immune compartment, dendritic cells, via diverse mechanisms of neuroimmune regulation involving neuropeptides, chemokines including CCL2 and electrical activity (Hanc et al. 2023).

Our study has further corroborated the remarkably pleiotropic and context-dependent effects of CGRP on diverse immune compartments across barrier tissues. In the skin, while sustained or potent activation of sensory neurons has been demonstrated to elicit (1) an anticipatory protective immune response in the context of fungal infection and optogenetic stimulation (Kashem et al. 2015; Cohen et al. 2019) or (2) an anti-inflammatory response to bacterial pathogens (Chiu et al. 2013; Pinho-Ribeiro et al. 2018), we propose that short-term or tonic releases of CGRP from sensory fibers in our setting of homeostatic host-microbiota interaction may moderate or constrain the adaptive immune response in the skin, possibly to circumvent collateral tissue damage or excessive metabolic cost. Notably, we observed that CGRP conferred pleiotropic effects on production of multiple cytokines beyond IL-17 by commensal-reactive T cells *ex vivo* (data not shown), further illustrating the versatile nature of CGRP as a neuroimmune modulator. The precise molecular mechanisms underlying how CGRP remodels the transcriptome and proteome of commensal-reactive T cells, which may in part involve PKA/cAMP response element-mediated signaling pathways, reminiscent of previous findings in the context of CGRP-dependent anti-inflammatory responses in dendritic cells and macrophages (Harzenetter et al. 2007; Altmayr, Jusek, and Holzmann 2010; Holzmann 2013), remain to be studied. Furthermore, CGRP may be able to modulate the immune system indirectly by reshaping the skin microbiota, potentially via a similar mechanism where substance P regulates the composition of the microbiota in the gut to promote tissue protection (Zhang et al. 2022).

Sources of physiological sensory modalities that can modify commensal-specific T cell activity and the tissue milieu may include temperature, noxious pain, mechanical stimuli, microbially derived molecules, as well as direct or indirect consequences of diets, skincare products, pain management or cancer treatment. Direct perturbations of systemic CGRP activity such as migraine treatment or prevention (Deen et al. 2017; Russo and Hay 2023) may also be able to impact commensal-specific lymphocytes at the barrier tissue, potentially contributing to side effects over the course of this powerful migraine therapeutic or prophylactic approach. Intriguingly, many side effects from migraine management that have been documented, such as exacerbated inflammation (Tracey 2002), altered course of wound healing or bruising (Wurthmann et al. 2020; Cullum et al. 2021) and alopecia (Ruiz et al. 2023), involved dysregulation at barrier tissues, raising an intriguing possibility that intervention with the CGRP- RAMP1 axis may have broader consequences at barrier sites in the clinical or human health context. With the emergence of acute migraine treatment approaches such as intranasal sprays (Reuter 2023), where CGRP modulatory molecules are directly deposited at barrier tissues such as the nasal mucosa, the uncharacterized impact of CGRP on local tissue physiology, whether adverse or beneficial to human health, warrants further studies.

The neuroimmune CGRP-RAMP1 signaling axis has been shown to influence lymphoid developmental stages beyond antigen-experienced T cells in the adult skin. Specifically, CGRP- RAMP1 promotes hematopoiesis and hematopoietic stem cell mobilization (Suekane et al. 2019; Gao et al. 2021), as well as thymocyte education (Kurz et al. 1995). Our characterization and re- contextualization of CGRP-RAMP1 signaling in commensal-induced T cells in the skin at homeostasis may reinforce the idea that the CGRP-RAMP1 axis is a temporally and spatially versatile rheostat of lymphocyte activity, rapidly tuning its activity to sensory modalities, particularly in commensal-specific T cells that are extremely plastic and are situated in the environment that constantly encounters changing physical and microbial cues. Because adaptive immune cell activation is already highly antigen-specific and antigen-dependent, CGRP may not directly mediate or modulate the antigen recognition steps. On the contrary, because the innate immune arm lacks specific antigen recognition, the input from the nervous system, such as CGRP, neuromedin and catecholamines, may play a more prominent role as a licensing agonist or an immunoregulatory factor in innate and innate-like immune cells (Klose et al. 2017; Moriyama et al. 2018; Yano and Artis 2022). We propose that CGRP participates in functional fine-tuning of commensal-specific adaptive immunity, which may have implications in shaping the dynamic interactions between innate and adaptive immune compartments.

It is notable that only the subset of adaptive lymphocytes involved in Type 17 responses, but not those mediating Type 1 responses, has evolved a CGRP-RAMP1-dependent mechanism to communicate directly with the sensory nervous system. Of note, cAMP-responsive elements downstream of the CGRP-RAMP1 axis have been shown to be involved in Th17 differentiation in the context of autoimmunity and inflammation (Hernandez et al. 2015; Yoshida et al. 2016). Because Type 17 responses are more associated with extracellular microbes and epithelial surfaces, we speculate that Tc17s and Th17s have evolved the ability to be tuned by both classical antigen recognition and neuronal signals, the latter of which are elicited by similar or relevant extracellular cues such as abiotic (temperature, noxious pain) and microbially derived (PAMPs/DAMPs) stimuli, which can in parallel directly activate sensory fibers (Yang and Chiu 2017) and trigger CGRP release. Such coordination between antigen-mediated signaling and neuroimmune tuning may allow RAMP1-expressing Type 17 commensal-specific T cells to respond more effectively to fluctuating cues at these epithelial surfaces. Of note, ILC3s, which regulate barrier integrity and defense through IL-17 and IL-22, are under the control of the VIP neuropeptide released from feeding-dependent enteric neurons (Talbot et al. 2020; Seillet et al. 2020; Yu et al. 2021), as well as neuromodulators produced by enteric glial cells (Ibiza et al. 2016), pointing to a paramount role of neuroimmune coordination in maintaining organ function and promoting barrier protection. Intriguingly, intestinal goblet cells, which play instrumental roles in maintaining barrier integrity, also respond to CGRP via RAMP1 to promote mucus production (Yang et al. 2022), illustrating another instance where sentinel cells at the epithelial surface is subject to regulation by this critical neuropeptide, which appears to mediate a wide range of multisystem crosstalk at barrier tissues.

Tuning of commensal-induced T cells by the neuroimmune CGRP-RAMP1 axis can functionally impact barrier tissues, as we showed that *Ramp1*-deficient T cells, refractory to the local neuronal signal and behaving in a manner that is less restrained in Type 17 immunity and activation status, were able to promote tissue repair or result in ectopic activation of keratinocytes. Because sustained, unfettered activation of T cells and concomitant hyperactivation of the epithelial tissue milieu may trigger collateral tissue damage, particularly in immunopathological settings, we propose that homeostatic CGRP-RAMP1-dependent tuning of commensal-reactive lymphocytes may serve as both a guardrail and a rheostat of adaptive immunity to moderate barrier tissue physiology, in the presence of ever-changing cues or potential threats from microbes and sensory modalities. In light of the ability of CGRP-RAMP1 signaling to modulate anti-tumor immunity mediated by CD8^+^ T cells (Balood et al. 2022), we anticipate that better understanding of how the neuroimmune CGRP-RAMP1 axis impacts homeostatic Type 17 adaptive immunity may contribute to a novel, tailored and rationalized approach to reprogram or customize T cells in various clinical settings, via harnessing neuromodulators to suppress or enhance adaptive immunity at the microbiota-colonized barrier surfaces in the appropriate context of human health and immunopathology.

## Materials and Methods

### Mice

Conventional Specific Pathogen Free (SPF) wild-type C57BL/6, *Foxp3-GFP* reporter (C57BL/6- Foxp3^tm1Kuch^) and Albino B6 (C57BL/6NTac-*Tyr^tm1Arte^*) mice were purchased from Taconic Bioscience. *Lck-Cre* (B6.Cg-Tg(Lck-icre)3779Nik/J and *Ox40-Cre* (B6.129X1(*Tnfrsf4^tm2(cre)Nik^*/J) mice were purchased from the Jackson Laboratory. *Ramp1 fl/fl* mice were provided by Dr. Isaac Chiu (Harvard Medical School), and were then bred at NIAID with the indicated *Cre* lines to generate *Cre-* and *Cre+* littermates in the *Ramp1 fl/fl* background. *Calca-EGFP* (Calca^tm1.1(EGFP/HBEGF)Mjz^) mice (McCoy, Taylor-Blake, and Zylka 2012) were kindly provided by Dr. John O’ Shea (National Institute of Arthritis and Musculoskeletal and Skin Diseases) with permission from the Mutant Mouse Resource & Research Centers (MMRRC). *Ramp1-/-* mice (B6.129S2(Cg)-*Ramp1^tm1.1Tsuj^*/WkinJ) were kindly provided by Dr. Wade Kingery (currently available at the Jackson Laboratory), and were then bred with the transgenic *S. epidermidis*-specific Bowie^Tg^ line (Harrison et al. 2019) to generate donor mice containing *Ramp1*-deficient GFP-expressing Bowie^Tg^ cells. Mice expressing OFP were kindly provided by Dr. Dorian McGavern (National Institute of Neurological Disorders and Stroke), and were then bred with the transgenic *S. epidermidis*-specific Bowie^Tg^ line to generate donor mice containing OFP-expressing Bowie^Tg^ cells. All mice were bred and maintained under specific pathogen-free conditions at an American Association for the Accreditation of Laboratory Animal Care (AAALAC)-accredited animal facility at NIAID and housed in accordance with the procedures outlined in the Guide for the Care and Use of Laboratory Animals. All experiments were performed under the animal study proposal LHIM-3E approved by the NIAID Animal Care and Use Committee. Unless otherwise noted, sex- and age-matched animals between 6 and 12 weeks of age were used for each experiment.

### Commensal bacterial culture and colonization

*Staphylococcus epidermidis* NIHLM087 was cultured for 18 hours in Tryptic Soy Broth at 37 °C without shaking. Up to 5ml of the bacterial suspension (an approximate density 10^9^ CFU/ml) was then applied topically to each mouse across the surface of the ear pinnae and/or the rest of the body (when indicated), using sterile cotton swabs. Topical association was performed every other day for four times for each experiment, where tissues from the animals were analyzed 14 days after the first commensal association. *Staphylococcus aureus* culture and colonization were described in Enamorado et al. 2023.

### Murine tissue processing

To obtain skin single-cell suspensions, ear pinnae were excised and separated into dorsal and ventral sheets, and then placed in RPMI 1640 media supplemented with 2 mM L-glutamine, 1 mM sodium pyruvate, 1 mM non-essential amino acids, 50 μM β-mercaptoethanol, 20 mM HEPES, 100 U/ml of penicillin, 100 mg/ml of streptomycin, 0.5 mg/ml of DNAse I (Sigma-Aldrich) and 0.25 mg/ml of Liberase TL purified enzyme blend (Roche) for 90 minutes at 37°C and 5% CO_2_. Digested ears were homogenized using the Medicon/Medimachine tissue homogenizer system (BD Biosciences) and filtered through 50-μm cell strainers.

### T cell in vitro restimulation

To assess cytokine production potential, single-cell suspensions were cultured in 96-well U-bottom plates for 2.5 h at 37°C in RPMI complete media (RPMI 1640 supplemented with 10% fetal bovine serum, 2 mM L-glutamine, 1 mM sodium pyruvate, 1 mM non-essential amino acids, 50 mM β-mercaptoethanol, 20 mM HEPES, 100 U/mL penicillin, and 100 mg/mL streptomycin) containing 50 ng/mL of phorbol myristate acetate (PMA; Sigma-Aldrich), 5 μg/mL ionomycin (Sigma-Aldrich), and a 1:1000 dilution of brefeldin A/GolgiPlug (BD Biosciences).

### Flow cytometry

Murine single-cell suspensions were incubated with fluorochrome-conjugated antibodies against surface markers and intracellular markers listed in **Supplemental Table S1**. Dead cells were excluded from live samples using 4′,6-diamidino-2-phenylindol (DAPI; Sigma-Aldrich) or LIVE/DEAD Fixable Blue Dead Cell Stain Kit (Invitrogen Life Technologies). For surface marker staining, cells were incubated with the indicated antibodies for 30 minutes at 4°C. For intracellular cytokine and transcription factor staining, cells were fixed and permeabilized using the Foxp3/Transcription Factor Staining Buffer Set (eBioscience) and stained with the indicated fluorophore-conjugated antibodies for at least 60 minutes at 4°C. All stainings were performed in the presence of purified anti-mouse CD16/32 (2.4G2, BioXcell). Data were acquired using BD Fortessa flow cytometers (BD Biosciences) running the FACSDiva software (BD Biosciences), and subsequently analyzed using FlowJo (v10, TreeStar).

### Ex vivo culture and cytokine production assessment

The indicated CD4^+^ and CD8^+^ T cell populations from the skin of commensal-colonized animals were isolated by FACS and cultured in the presence of TCR stimulation (1 μg/ml plate-bound anti-CD3e mAb, clone 145-2C11). Rat CGRP (α isoform; Sigma-Aldrich) at the indicated concentrations was added to the culture, and supernatants were collected 24 hours later and subjected to the FlowCytomix bead array assay (eBioscience) for cytokine production assessment.

### Confocal microscopy

For whole-mount visualization, mouse ears were excised and separated into ventral and dorsal sheets using fine forceps and the microscope. Tissues were fixed in 4% paraformaldehyde solution (Electron Microscopy Sciences) overnight at 4°C with shaking. Fixed tissues were then blocked with 1% BSA, 0.25% Triton X-100 and Fc Block for 2 hours at room temperature with shaking. Tissues were first stained with the indicated fluorophore-conjugated or primary antibodies in blocking solution overnight at 4 °C with gentle shaking, washed with PBS three times at room temperature with shaking. When applicable, secondary antibody was performed for 60 minutes at room temperature followed by washing. After completion of staining and washing, tissues were mounted with ProLong Gold (Invitrogen) antifade reagent with the dermis facing the coverslip. After drying for 16 hours, images were captured with a Leica TCS SP8 confocal microscope equipped with HyD and PMT detectors and an HC PL APO 40×/1.3 oil objective. Images were analyzed using Imaris software (Bitplane).

### Intravital microscopy and analyses

An albino version of the *Calca-EGFP* reporter was generated by breeding albino B6 animals with *Calca-EGFP* animals. Albino mice heterozygous for *Calca-EGFP* were used for intravital imaging after an adoptive transfer with fluorochrome-expressing Bowie^Tg^ cells (approximately 5 x 10^5^ cells), followed by the same course of topical association with *S. epidermidis* (NIHLM087) described above. Intravital imaging was done 8-12 days after the initial commensal colonization to ensure optimal accumulation of fluorochrome-expressing Bowie^Tg^ cells in the ear skin for visualization.

Prior to intravital imaging, mice were injected with 25 µg of Alexa Fluor-647 labeled CD31 antibody (MEC13.3, BioLegend) retro-orbitally, in a total volume of 50 µL to enable visualization of the endothelium. Intravital multiphoton microscopy was performed using Leica Mi8 DIVE (Deep In Vivo Explorer) inverted confocal microscope (Leica Microsystems) equipped with dual multiphoton lasers (Spectra Physics). Mai Tai DS was used for excitation of the EGFP signal, and InSight DS for red and far-red probes. The microscope was additionally equipped with 4 ultra-sensitive HyD detectors, L 25.0 water-immersion objective (0.95 NA), a motorized stage, and Environmental Chamber (NIH Division of Scientific Equipment and Instrumentation Services) to maintain 37 °C for anesthetized animals. Mai Tai was tuned to 880 nm excitation, and InSight to 1150 nm excitation wavelengths. For non-invasive time-lapse imaging of the ear pinnae, tiled images of 2x2 fields were defined using Tilescan application of Leica Application Suite X (LAS X), and Z stacks consisting of 3-5 single planes (5-7 μm each over a total tissue depth of 30–50 μm) were acquired every 45 seconds for a total observation time between 1 to 6 hours. Raw imaging data were processed using Imaris (Bitplane). All imaging files were stabilized and adjusted for drifts prior to subsequent analysis. Cells (Bowie^Tg^) were surface-rendered using the Imaris Surface module to generate 3-D positional and morphological data at all time points. Skin sensory nerves (expressing *Calca-EGFP*) and endothelia (CD31^+^) were filament-rendered using the Imaris Filament module, and then surface-rendered for all time points. The distances between the rendered T cell surfaces and the rendered nerve and endothelium surfaces were calculated using shortest distance (object-object) calculation module, and the data from all time points from all the experimental animals were collected and analyzed.

### Single-cell RNA-sequencing and analysis

Skin CD8^+^ lymphocytes labeled with TotalSeqC hashtag antibodies (BioLegend) from *S. epidermidis*-associated mice (six *Lck-Cre- Ramp1 fl/fl* animals and six *Lck Cre+ Ramp1 fl/fl* animals) were sorted as DAPI- CD90.2^+^ TCRβ^+^ CD4^-^ CD8β^+^ TCRγδ^-^ CD49f^-^ NK1.1^-^ B220^-^ CD11b^-^ CD11c^-^ MHCII^-^, using a Sony MA900 cell sorter. All samples were pooled together, and approximately 15,000 sorted cells were loaded onto the Chromium Single Cell Controller (10X Genomics) to encapsulate cells into droplets. The library was prepared using a Chromium Single Cell 5’ Reagent Kits v2 (10X Genomics) following manufacturer’s instructions, and then sequenced on an Illumina Nextseq500 (Next Seq 500/550 High Output Kit v2, Illumina). Data were filtered and mapped to mm10 reference genome using cellranger 7.0.0 (10X Genomics). Data were normalized in Seurat and were displayed as uniform manifold approximation and projection (UMAP). Three out of thirteen Seurat clusters were excluded from the analyses shown, as each of those outlier clusters only represented zero to ten cells (less than one percent) per cluster per animal. Differential gene expression for each cluster was performed using the Seurat FindAllMarkers function with a single cell-tailored test MAST and default parameters. Gene Ontology (GO) enrichment analysis for differentially expressed genes were performed using Metascape (Zhou et al. 2019). The input for GO enrichment analysis were DEGs with p-values less than 0.05 from the indicated Seurat clusters.

Furthermore, publicly available scRNA-seq datasets, previously generated in our laboratory, were used for new analyses. For **Figures 2D, 2E** and **2F,** the scRNA-seq dataset from sorted commensal-induced CD8^+^ T cells from Harrison et al., 2019 was used for analyses (NCBI BioProject PRJNA486019). For **Figure S2F,** the scRNA-seq dataset from sorted MAIT cells from Constantinides et al., 2019 was used for analyses (NCBI BioProject PRJNA529261).

### Bulk RNA-sequencing and analysis

CD4^+^ T cells from wild-type animals following *S. epidermidis* colonization were sorted into Th1 and Th17, processed and analyzed for RNA-seq as described in Linehan et al., 2018. Keratinocytes from the same *Lck-Cre Ramp1 fl/fl* experiment described above were sorted and processed for RNA-seq as described in Lima-Junior et al., 2021. Sequencing reads were mapped to the mm10 (GRCm38) mouse genome using STAR with stringent mapping conditions, where reads aligned to multiple regions of the genome were eliminated (-outFilterMultimapNmax 2). Differential gene expression was calculated using HOMER’s getDiffExpression (Heinz et al. 2010). Genes with FDR < 0.05 and foldchange > 2 were considered differentially expressed transcripts. Gene Ontology (GO) enrichment analysis for differentially expressed genes was performed using Metascape (Zhou et al. 2019). Retroelement and noncoding transcript expression was determined using HOMER 4.11 (Heinz et al. 2010), defining these transcripts based on the annotations provided by the Genome-based Endogenous Viral Element Database (gEVE) for GRCm38. For the analysis of individual loci, reads were analyzed using HOMER’s analyzeRepeats function (Heinz et al. 2010), along with a custom GFT file with the gEVE annotations. For the analysis of retroelement and noncoding transcript family motifs, reads were analyzed using the analyzeRepeats function with parameters “repeats mm10”.

Furthermore, a publicly available RNA-seq dataset, previously generated in our laboratory, from sorted CD8^+^ T cells under various conditions from Linehan et al., 2018 was used for analyses in **Figures 2A, 2B, 2C, 2G, S2A, S2B, S2C** and **S2D**. The dataset is available on NCBI BioProject PRJNA419368. Additionally, a publicly available RNA-seq dataset, previously generated in our laboratory, from sorted skin CD4^+^ T cells after *S. aureus* topical association from Enamorado et al., 2023 was used for analyses in **Figure 2G**. The dataset is available on NCBI GEO accession GSE196994.

### ATAC-sequencing

A publicly available ATAC-seq dataset, previously generated in our laboratory, from sorted CD4^+^ and CD8^+^ T cells under various conditions from Harrison et al., 2019 was used for analyses in **Figures 2H, S2G, 3H, S3C** and **S3D**. The dataset is available on NCBI BioProject PRJNA486019.

To scan for motif occurrences, HOMER’s scanMoifGenomeWide.pl was used (Heinz et al. 2010). The results were visualized in the UCSC Genome Browser (Kent et al. 2002) by uploading custom hubs.

### Wounding and epifluorescence microscopy of wound tissue

*S. epidermidis*-associated male mice in the telogen phase of the hair cycle were anesthetized and shaved. A 6-mm biopsy punch was utilized to partially perforate the skin, and iris scissors were then used to cut the skin along the punch outline. Five days after wounding, the skin tissue was excised and fixed in 4% paraformaldehyde for 4 hours at 4°C, incubated in 30% sucrose at 4°C overnight, embedded in OCT compound (Tissue-Tek), frozen on dry ice, and cryo-sectioned (20- μm-thick sections). Sections were then fixed in 4% paraformaldehyde at room temperature for 10 minutes, permeabilized with 0.1% Triton X-100 (Sigma-Aldrich) for 10 minutes, and blocked for 1 hour at room temperature in blocking buffer (2.5% normal goat serum, 1% BSA, 0.3% Triton X-100 in PBS). Sections were stained with a chicken anti-mouse Keratin 14 antibody (Poly9060, BioLegend), diluted at 1:400 in blocking buffer containing rat gamma globulin and anti- CD16/32, with an overnight incubation at 4°C. After washing with PBS, sections were then stained with a polyclonal anti-chicken IgY-AlexaFluor647 secondary antibody (Jackson ImmunoResearch) diluted at 1:800 at room temperature for 1 hour, stained with DAPI and mounted with ProLong Gold (Invitrogen). Wound images were captured with a Leica DMI 6000 widefield epifluorescence microscope equipped with a Leica DFC360X monochrome camera. Tiled and stitched images of wounds were collected using a 20X/0.4 NA dry objective. Images were analyzed using Imaris (Bitplane) and quantitation was done in a double-blind manner.

### Statistical analysis

Prism 9 (GraphPad) was used to determine the indicated statistical significance.

## Acknowledgments

W.K. was supported by the Damon Runyon Cancer Research Foundation. Y.B. is supported by the Division of Intramural Research of NIAID (1ZIA-AI001115 and 1ZIA-AI001132). We thank Dr. Margery Smelkinson, Galina Koroleva, Dr. Alexander T. Chesler, Dr. Seong-Ji Han, Kimberly Beacht, Ejae Lewis, Dr. Juliana Perez-Chaparro and Brittany Dulek for technical support, advice and assistance with experiments. We thank Dr. Dorian McGavern, Dr. John O’Shea and Dr. Wade Kingery for generously sharing the indicated mouse lines. We thank members of the Belkaid lab for technical input and assistance, as well as constructive feedback throughout the research project and the writing of the manuscript.

---

**Figure S1.**
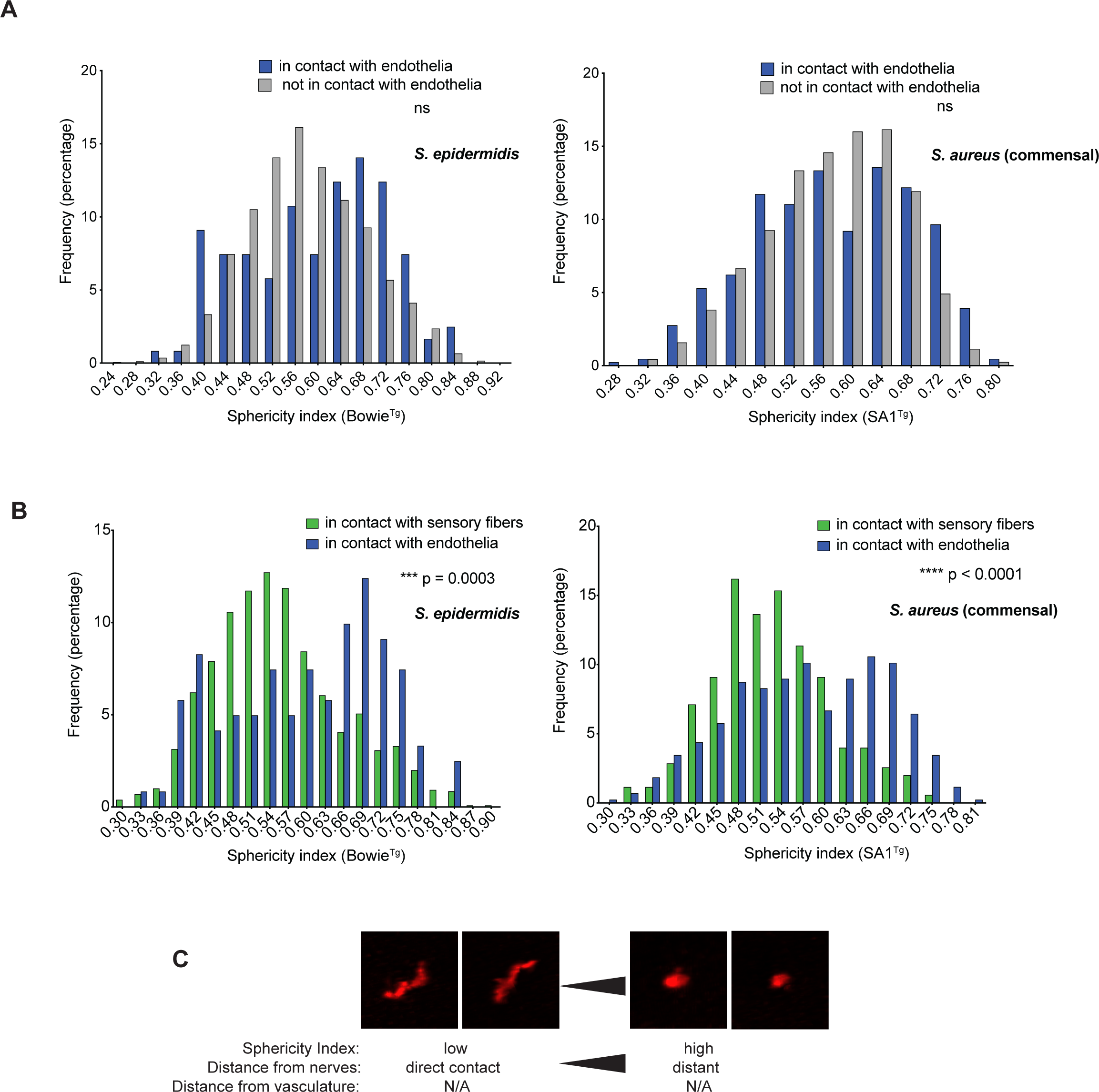
Proximity to sensory nerves impacts commensal-specific T cell behavior at homeostasis *in vivo* **(A)** Each graph represents distribution of cellular sphericity values of commensal-specific T cells (Bowie^Tg^ for *S. epidermidis* and SA1^Tg^ for *S. aureus*) over the recording period. Bars in the indicated colors represent populations of the T cells that are in direct contact or not in direct contact with the endothelium. **(B)** Each graph represents distribution of cellular sphericity values of commensal-specific T cells (Bowie^Tg^ for *S. epidermidis* and SA1^Tg^ for *S. aureus*) over the recording period. Bars in the indicated colors represent populations of the T cells that are in direct contact or not in direct contact with the indicated structures. **(C)** A schematic summarizing the relationship between distances from nerve fibers or the vasculature and cellular shape *in vivo* over the recording period. All data shown in (**A**)-(**B**) represent at least three independent experiments. P-values were calculated with Student’s t test. “ns” denotes not statistically significant.

**Figure S2.**
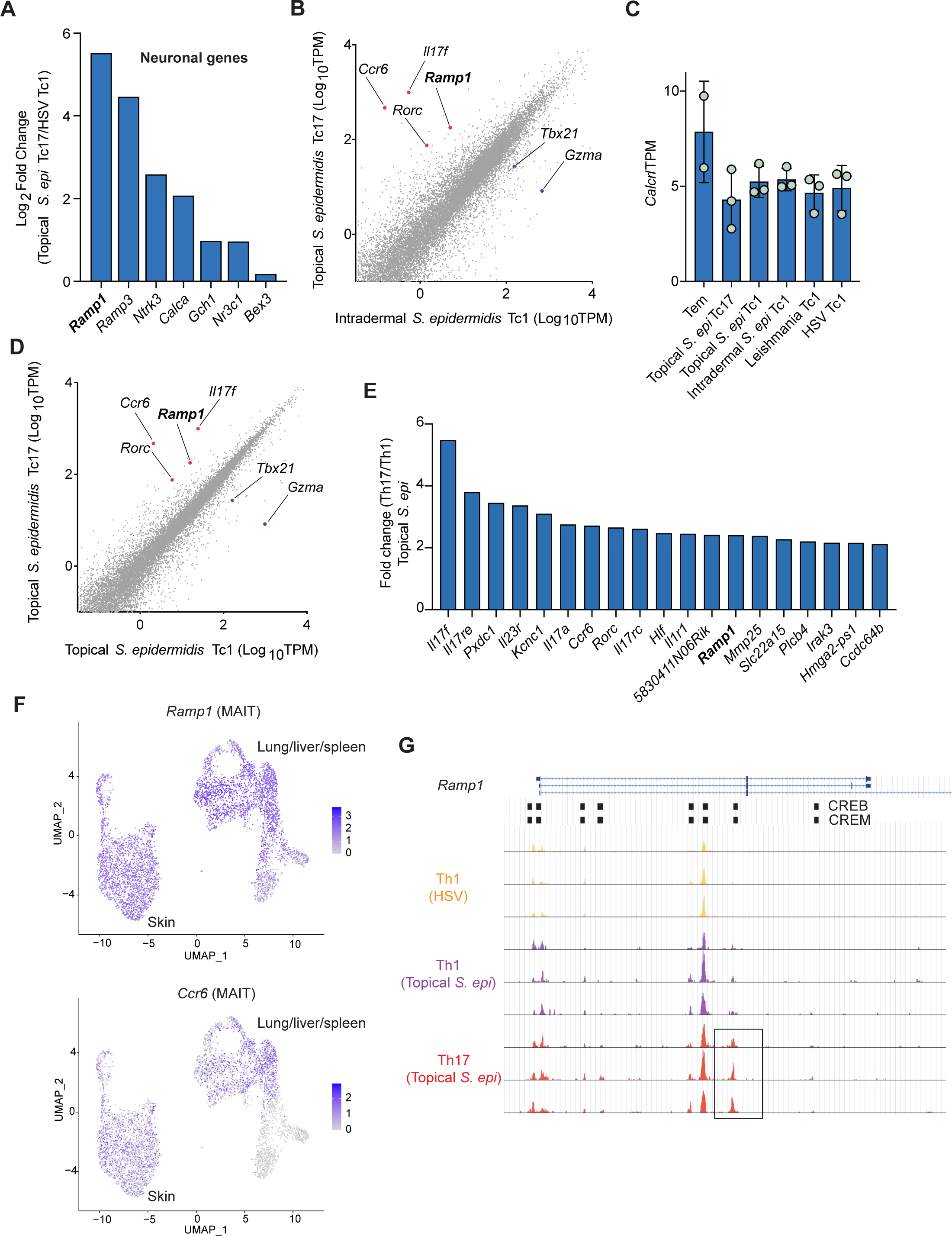
Expression of the gene encoding the CGRP receptor, *Ramp1*, is enriched in Type 17 commensal-induced T cells *in vivo* **(A)** Bar graph showing upregulation of the indicated neuronal genes in commensal-induced Tc17 cells, compared to Tc1 cells elicited by HSV infection. **(B)** Scatter plot showing differentially expressed genes, comparing topical *S. epidermis*- induced Tc17 cells vs intradermal (infection) *S. epidermis*-induced Tc1 cells from the skin. Red and blue dots highlight differentially expressed genes of interest. **(C)** Bar graph showing expression levels of *Calcrl*, encoding a co-receptor for CGRP, in the indicated CD8^+^ T cell populations. **(D)** Scatter plot showing differentially expressed genes, comparing topical *S. epidermis*- induced Tc17 cells vs Tc1 cells from the skin. Red and blue dots highlight differentially expressed genes of interest. **(E)** Bar graph showing upregulation of *Ramp1* and other Type 17 response genes in commensal-induced Th17 cells compared to Th1 cells elicited by *S. epidermidis* topical colonization. **(F)** Feature plots showing expression of *Ramp1* and *Ccr6* in MAIT cells from different barrier tissues. **(G)** UCSC Genome Browser image of open chromatin regions of the *Ramp1* locus in the indicated CD4^+^ T cell populations from the ATAC-seq data. Refer to detailed RNA-seq and ATAC-seq data sources and experimental settings in Figure 2 and **Materials and Methods**. Bar graphs show mean ± standard deviation.

**Figure S3.**
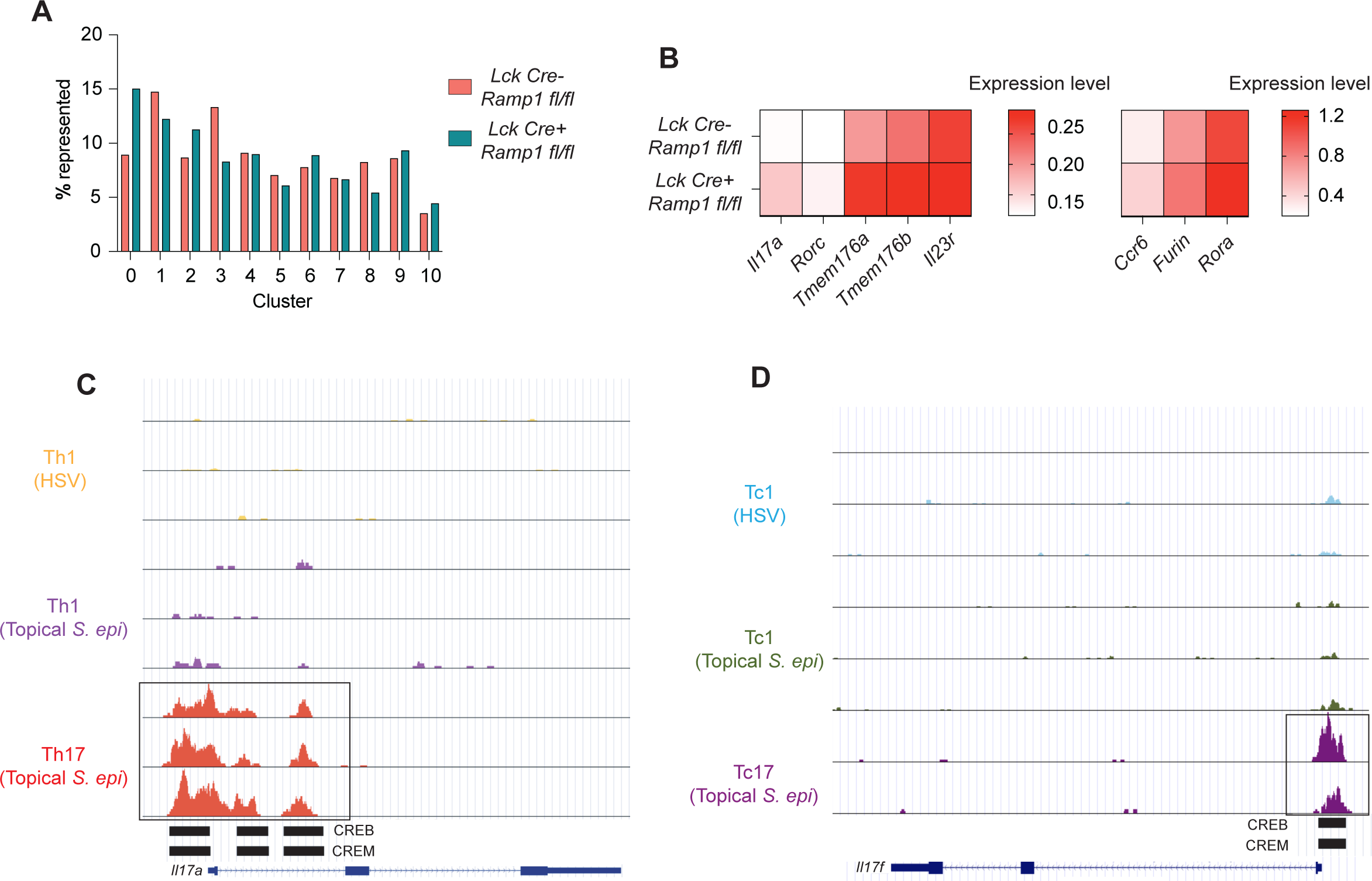
Neuroimmune CGRP-RAMP1 signaling constrains Type 17 responses in commensal-induced T cells *in vivo* **(A)** Graph showing the frequency of cell numbers from each genotype (*Ramp1*-deficient vs control genetic background) for each assigned Seurat cluster. Data from the scRNA-seq experiment described in Figure 3. **(B)** Heatmap showing the expression levels of the indicated Type 17 genes for each genotype (*Ramp1*-deficient vs control genetic background). Data from the scRNA-seq experiment described in Figure 3. **(C)** UCSC Genome Browser image of open chromatin regions of the *Il17a* locus in the indicated CD4^+^ T cell populations in the wild-type genetic background from the ATAC-seq data described in Figure 2. **(D)** UCSC Genome Browser image of open chromatin regions of another Type 17 response gene, *Il17f*, locus in the indicated CD8^+^ T cell populations in the wild-type genetic background from the ATAC-seq data described in Figure 2.

**Figure S4.**
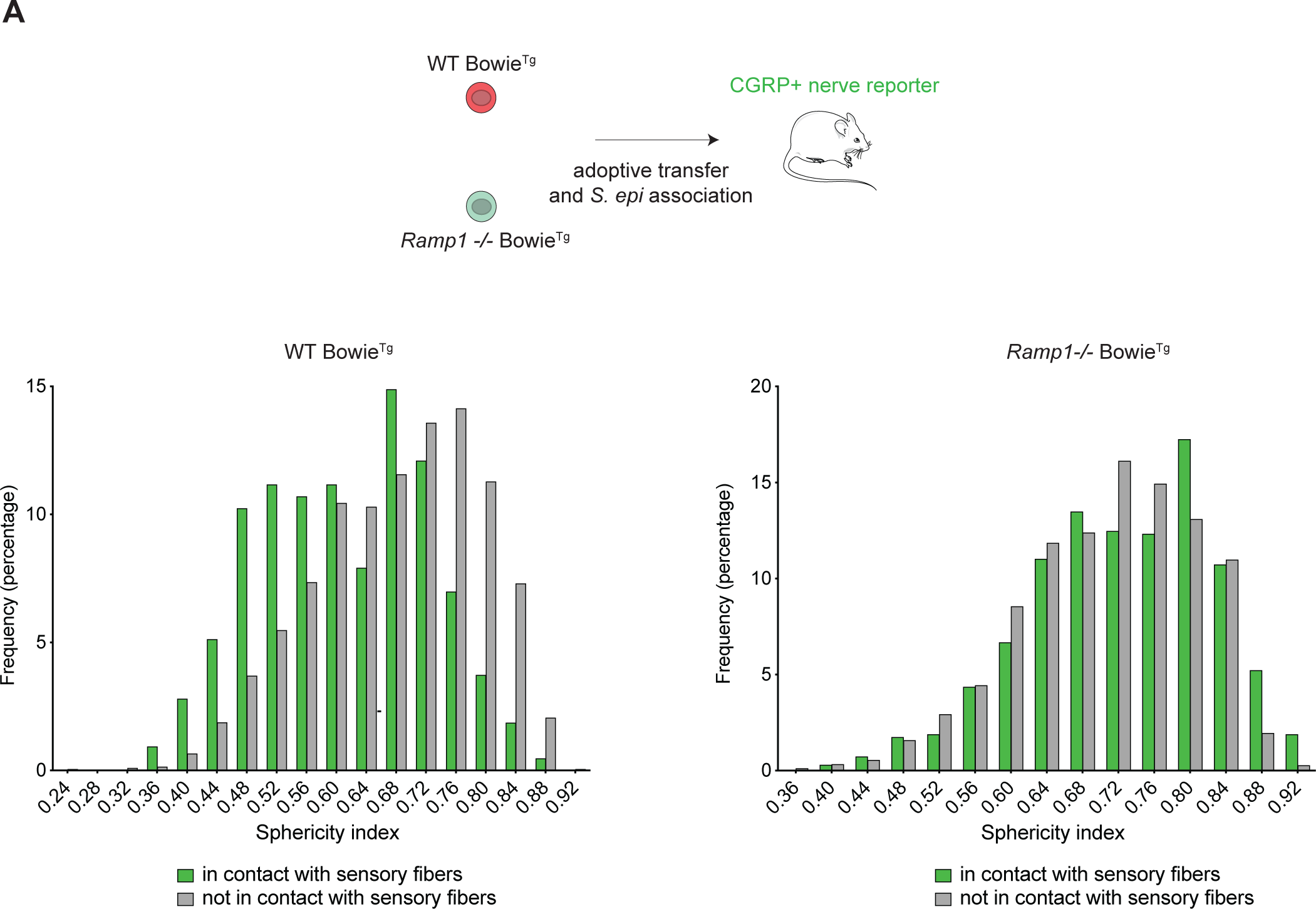
The neuroimmune CGRP-RAMP1 axis controls the ability of sensory nerve fibers to impact commensal-specific T cell behavior *in vivo* **(A)** Each graph represents distribution of cellular sphericity indexes of commensal-specific T cells over the recording period. Bars in the indicated colors represent populations of the T cells that are in direct contact or not in direct contact with sensory fibers. For the wild-type commensal-specific T cells, the mean sphericity index difference between the populations that were in direct contact vs not in direct contact was 0.07 (**** p < 0.0001), whereas such mean sphericity index difference was 0.02 (** p = 0.004) for the *Ramp1*-deficient commensal-specific T cells.

**Figure S5.**
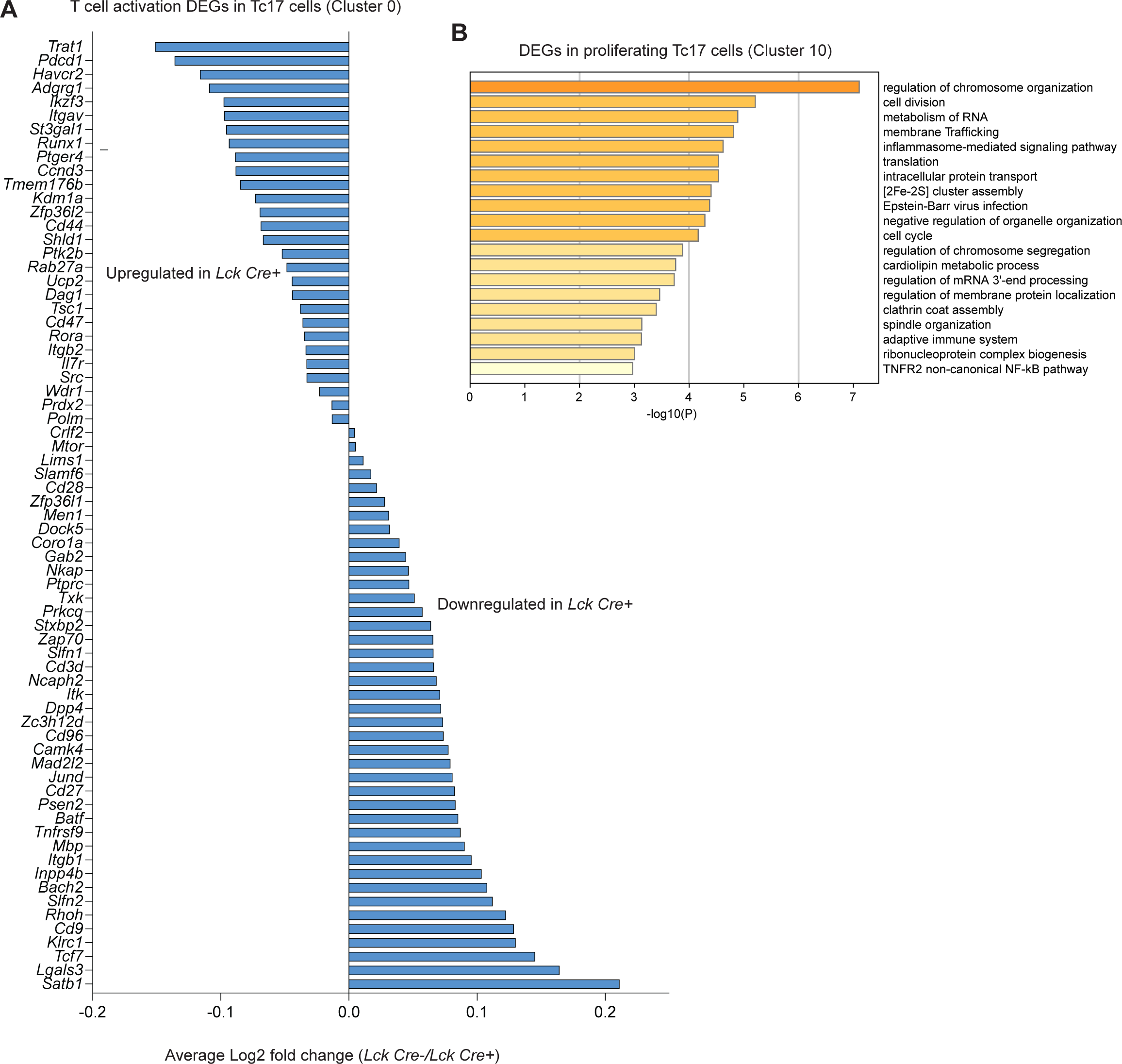
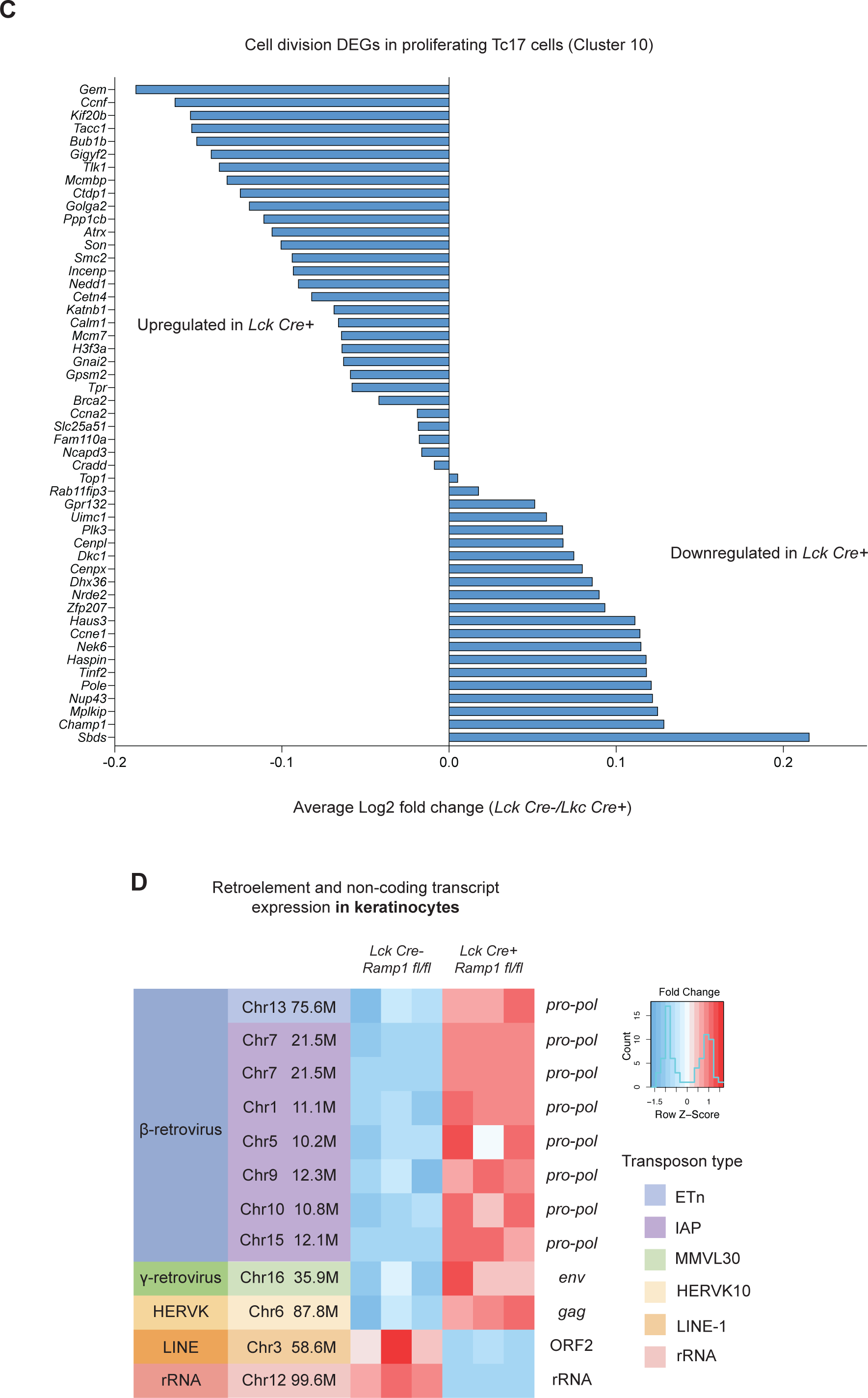
Neuroimmune CGRP-RAMP1 signaling modulates expression levels of activation and proliferation genes in commensal-induced Tc17 populations **(A)** Fold changes of differentially expressed genes involved in T cell activation in the dominant non-proliferating Tc17 cluster (Cluster 0 from Figure 3), based on the GO enrichment analysis in Figure 4B. **(B)** GO enrichment analysis of differentially expressed genes in the proliferating Tc17 cluster (Cluster 10 from the scRNA-seq dataset described in Figure 3), comparing commensal-induced T cells from *Ramp1*-deficient vs control genetic backgrounds. **(C)** Fold changes of differentially expressed genes involved in cell division in the proliferating Tc17 cluster (Cluster 10 from Figure 3), based on the GO enrichment analysis in **Figure S5B**. **(D)** Analysis of differentially expressed retroelement and non-coding transcript loci, comparing keratinocytes isolated from the tissue milieu where commensal-induced T cells were *Ramp1*-deficient vs control.

Supplemental Video 1. Commensal-specific T lymphocytes are in close proximity to skin sensory nerve fibers at homeostasis *in vivo*

Videos showing dynamic interactions between commensal-specific T cells and the indicated neural or endothelial structures *in vivo*. All three videos illustrate the same region and recording period, with different visual markers highlighted, representative of at least three independent experiments. Videos can be played synchronously under presentation mode.

Supplemental Table 1. Reagents

