## Supplemental Table 1 for "The neuroimmune CGRP-RAMP1 axis tunes cutaneous adaptive immunity to the microbiota"

| REAGENT or RESOURCE | SOURCE | IDENTIFIER | |
| --- | --- | --- | --- |
| Antibodies | | | |
| Anti-mouse CCR6, PE (29-2L17) | Biolegend | RRID: AB_1279137 | Cat# 129804 |
| Anti-mouse CD4, BV510 (RM4-5) | Biolegend | RRID: AB_2562608 | Cat# 100559 |
| Anti-mouse CD8β, BV650 (eBioH35-17.2) | eBioscience | RRID: AB_2921045 | Cat# 416-0081-82 |
| Anti-mouse CD8β, PE-Cy7 (eBioH35-17.2) | eBioscience | RRID: AB_11218494 | Cat# 25-0083-82 |
| Anti-mouse CD11b, eFluor 450 (M1/70) | eBioscience | RRID: AB_1582236 | Cat# 48-0112-82 |
| Anti-mouse CD11c, eFluor 450 (N418) | eBioscience | RRID: AB_1548654 | Cat# 48-0114-82 |
| Anti-mouse CD31, PerCP-Cy5.5 (MEC13.3) | Biolegend | RRID: AB_25566761 | Cat# 102522 |
| Anti-mouse CD34, eFluor 660 (RAM34) | eBioscience | RRID: AB_10596826 | Cat# 50-0341-82 |
| Anti-mouse CD45, APC-eFluor 780 (30-F11) | eBioscience | RRID: AB_1548781 | Cat# 47-0451-82 |
| Anti-mouse CD49f, eFluor 450 (eBioGoH3) | eBioscience | RRID: AB_11042564 | Cat# 48-0495-82 |
| Anti-mouse CD49f, PE (eBioGoH3) | eBioscience | RRID: AB_893373 | Cat# 313612 |
| Anti-mouse CD90.2, BV785 (30-H12) | Biolegend | RRID: AB_2562900 | Cat# 105331 |
| Anti-mouse MHC-II, eFluor 450 (M5/114.15.2) | eBioscience | RRID: AB_1272204 | Cat# 48-5321-82 |
| Anti-mouse NK1.1, eFluor 450 (PK136) | eBioscience | RRID: AB_2043877 | Cat# 48-5941-82 |
| Anti-mouse Sca-1, FITC (D7) | Biolegend | RRID: AB_313343 | Cat# 108106 |
| Anti-mouse TCRβ, PerCP-Cy5.5 (H57-597) | eBioscience | RRID: AB_925763 | Cat# 45-5961-82 |
| Anti-mouse TCRβ, BUV737 (H57-597) | eBioscience | RRID: AB_2870145 | Cat# 612821 |
| Anti-mouse TCRγδ, eFluor 450 (eBioGL3) | eBioscience | RRID: AB_2574071 | Cat# 48-5711-82 |
| Anti-mouse TCRγδ, PE-CF594 (eBioGL3) | eBioscience | RRID: AB_2661844 | Cat# 563532 |
| Anti-mouse IFN-γ, BV605 (XMG1.2) | eBioscience | N/A | Cat# 406-7311-82 |
| Anti-mouse IL-17A, PECy7 (TC11-18H10.1) | Biolegend | RRID: AB_2125010 | Cat# 506922 |
| Anti-mouse IL-17F, AF488 (O79-289) | eBioscience | RRID: AB_10715832 | Cat# 561631 |
| Anti-mouse alpha-CGRP | Peninsula laboratories | RRID: AB_518147 | Cat# T-4032 |
| Normal Goat Serum | Jackson ImmunoResearch Laboratories | RRID: AB_2336990 | Cat #005-000-121 |
| Rat Gamma Globulin | Jackson ImmunoResearch Laboratories | RRID: AB_2337135 | Cat #012-000-002 |
| Normal rabbit Serum | Jackson ImmunoResearch Laboratories | RRID: AB_2337123 | Cat #011-000-120 |
| Bacterial and Virus Strains | | | |
| *Staphylococcus epidermidis* NIHLM087 | Laboratory of Dr. Julie Segre (NHGRI/NIH) | N/A | |
| Chemicals, Peptides, and Recombinant Proteins | | | |
| 2-Mercaptoethanol (1,000X) | Gibco | Cat # 21985-023 | |
| 2-Mercaptoethanol | Sigma-Aldrich | Cat # M3148-25ML | |
| BSA | Sigma-Aldrich | Cat #A3059-500G | |
| Brefeldin A (GolgiPlug) | BD Biosciences | Cat# 555029 | |
| DAPI | Sigma-Aldrich | Cat # D9542 | |
| DMEM medium | Corning | Cat # 10-017-CV | |
| DNase I | Sigma-Aldrich | Cat# DN25-5G | |
| EDTA (0.5M) | Corning | Cat # 46-034-Cl | |
| FBS | Hyclone | Cat # SH30070.03 | |
| L-Glutamine | Corning | Cat # 25-005-Cl | |
| HEPES | Corning | Cat # 25-060-Cl | |
| Ionomycin | Sigma-Aldrich | Cat # I0634-5MG | |
| Liberase TL | Roche | Cat# 05401020001 | |
| MEM Non-essential Amino Acids (100X) | Corning | Cat # 25-025-Cl | |
| Paraformaldehyde | Electron Microscopy Sciences | Cat # 15714-S | |
| Pennicillin-Streptomycin (100X) | Corning | Cat # 30-002-Cl | |
| Phorbol 12-myristate 13-acetate (PMA) | Sigma-Aldrich | Cat # P8139-10MG | |
| ProLong Gold Antifade Mountant | Molecular Probes | Cat # P36930 | |
| RNAlater | Sigma-Aldrich | Cat # R0901-100ML | |
| RPMI 1640 medium | Corning | Cat # 10-040-CV | |
| Sodium Pyruvate (100X) | Corning | Cat # 25-000-Cl | |
| Triton X | Sigma-Aldrich | Cat #T9284 | |
| Critical Commercial Assays | | | |
| BD Cytofix/Cytoperm | BD Biosciences | Cat# 554722 | |
| BD Perm/Wash | BD Biosciences | Cat# 554723 | |
| Foxp3 / Transcription Factor Staining Buffer Set | eBioscience | Cat# 00-5523-00 | |
| High Sensitivity D1000 ScreenTape | Agilent | Cat # 5067-5584 | |
| LIVE/DEAD Fixable Blue Dead Cell Staining Kit | Life Technologies | Cat# L23105 | |
| MACS Cell Separation Column LS | Miltenyi Biotec | Cat #130-042-401 | |
| NextSeq 500/550 v2 kits (75 cycles) | Illumina | Cat# FC-404-2005 | |
| Qubit dsDNA HS Assay Kit | Molecular Probes | Cat # Q32854 | |
