## Supplemental Video 1 for "The neuroimmune CGRP-RAMP1 axis tunes cutaneous adaptive immunity to the microbiota"

### Slide 1
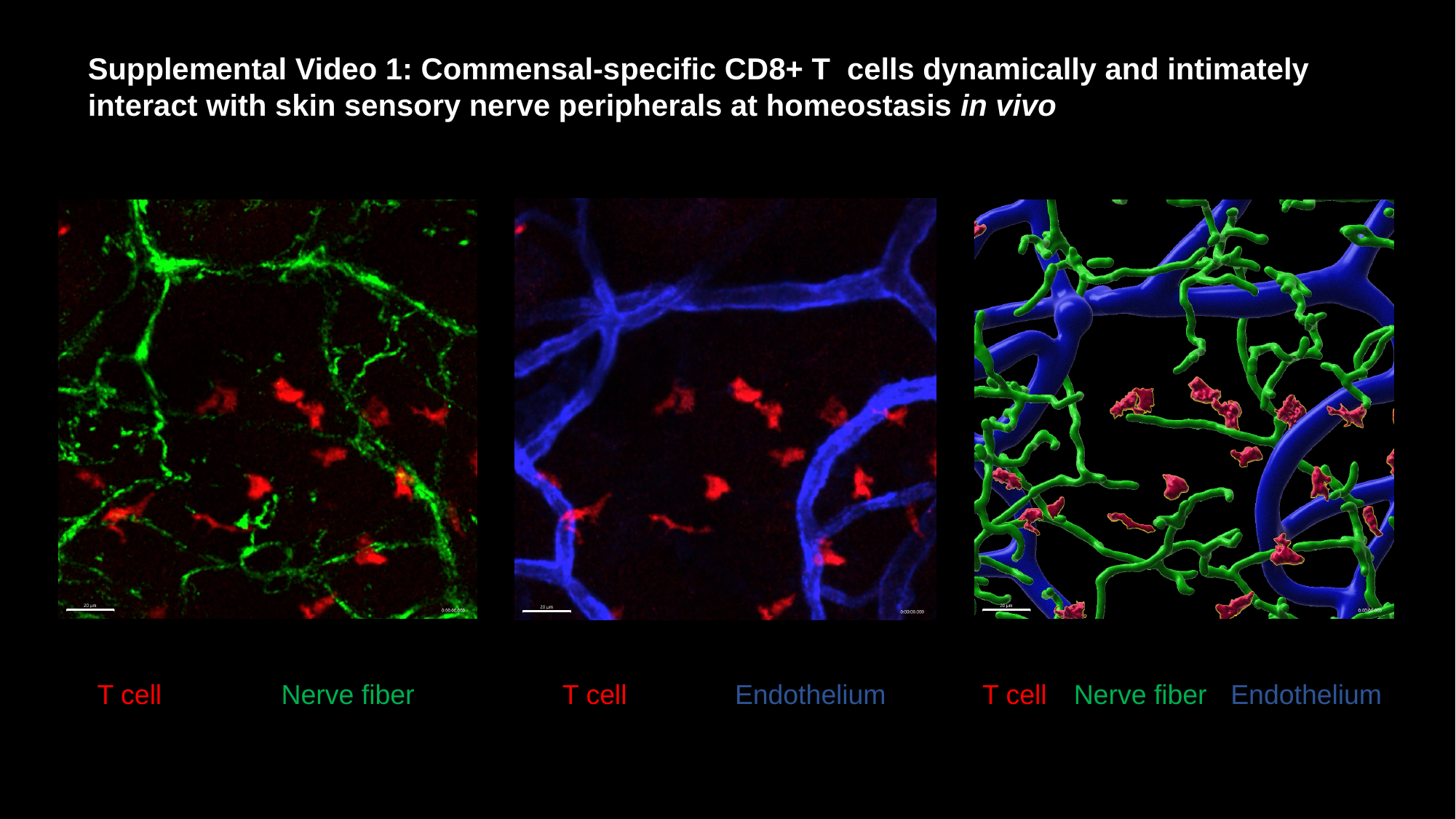

Supplemental Video 1: Commensal-specific CD8+ T cells dynamically and intimately interact with skin sensory nerve peripherals at homeostasis in vivo
T cell
Nerve fiber
T cell
Endothelium
T cell
Nerve fiber
Endothelium
